# Leveraging non-human primate multisensory neurons and circuits in assessing consciousness theory

**DOI:** 10.1101/584516

**Authors:** Jean-Paul Noel, Yumiko Ishizawa, Shaun R. Patel, Emad N. Eskandar, Mark T. Wallace

## Abstract

Both the Global Neuronal Workspace (GNW) and Integrated Information Theory (IIT) posit that highly complex and interconnected networks engender perceptual awareness. GNW specifies that activity recruiting fronto-parietal networks will elicit a subjective experience, while IIT is more concerned with the functional architecture of networks than with activity within it. Here, we argue that according to IIT mathematics, circuits converging on integrative vs. convergent yet non-integrative neurons should support a greater degree of consciousness. We test this hypothesis by analyzing a dataset of neuronal responses collected simultaneously from primary somatosensory cortex (S1) and ventral premotor cortex (vPM) in non-human primates presented with auditory, tactile, and audio-tactile stimuli as they are progressively anesthetized with Propofol. We first describe the multisensory (audio-tactile) characteristics of S1 and vPM neurons (mean and dispersion tendencies, as well as noise-correlations), and functionally label these neurons as convergent or integrative according to their spiking responses. Then, we characterize how these different pools of neurons behave as a function of consciousness. At odds with the IIT mathematics, results suggest that convergent neurons more readily exhibit properties of consciousness (neural complexity and noise correlation) and are more impacted during the loss of consciousness than integrative neurons. Lastly, we provide support for the GNW by showing that neural ignition (i.e., same trial co-activation of S1 and vPM) was more frequent in conscious than unconscious states. Overall, we contrast GNW and IIT within the same single-unit activity dataset, and support the GNW.

## Introduction

Understanding the neural architecture enabling arousal or wakefulness (i.e., level or state of consciousness) and conscious experience (i.e., content of consciousness) remains a central unanswered question in systems neuroscience despite its profound clinical implications in coma, vegetative-state, minimal-consciousness, and general anesthesia [1-3]. While in recent years a number of electrophysiological measures of consciousness/awareness have been proposed [4-6], these tend to be more practical than principled, and grounded more in the realm of engineering than neurobiology.

Lacking a mechanistic account of consciousness, a number of theorists and researchers have started from empirical observations or phenomenological axioms to derive consciousness theories. A number of these theories share many commonalities - as well as a number of practical and conceptual differences – as exemplified by two of the most prevailing and influential of these theories: Global Neuronal Workspace (GNW; [7, 8]) and Integrated Information Theory (IIT;[9-11], but see [12-17], for a number of other theories).

The GNW posits that an external stimulus will evoke a conscious experience if the associated neural information is widely distributed across distinct brain areas and networks – most prominently in the pre-frontal cortex [8]. Rendering sensory information globally available results in a coherent neural assembly of sustained activity and is most readily indexed via “neural ignition”, the non-linear process whereby in unconscious states neuronal activation profiles remain encapsulated within their specialized subsystems, whereas in conscious experiences these activation patterns are widely distributed (see [18] for experimental evidence and [19] for a recent computational treatise).

In a similar manner, IIT is a systems-level theory of consciousness also postulating that complex and highly interconnected neural networks support subjective experience [11, 20]. In contrast to GNW, however, the mathematics developed in hand with the IIT [9, 11] are argued to apply to all physical networks, and in turn IIT is arguably more focused on the architecture of networks rather than the activity within these. In more detail, Tononi and colleagues argue that each conscious experience is highly informative, as it represents a particular instance among a vast repertoire of potential experiences, and is highly unified, unable of being deconstructed into sub-experiences that are each independently perceived [21]. In turn, the IIT specifies that an organism may support conscious experience if imbued with an information processing architecture that is capable of supporting both high differentiation (i.e., large repertoire of possible states) and integration (i.e., strong statistical dependencies between system components).

Unfortunately, the neurophysiological data that bears directly on these theories is limited, in particular the IIT given its computational overhead. Indeed, a strength of the IIT is that it explicitly generates a metric of consciousness level; phi (Φ). This value, phi, can in principle be computed for any information processing system, as long as the transition probability matrix between nodes of the system are known, and in essence is proportional to the amount of information gained by knowing the state of all nodes within the system vs. having access to a limited purview of the system (see [9] for more detail). Regretably, computing this measure in complex biological systems is impossible from a practical standpoint due to its combinatorial search problem (but see [22, 23] for interesting approaches circumventing current computing limitations).

In an effort to provide empirical evidence germane to theories of consciousness, we propose here simple neurophysiological benchmarks for consciousness as derived from the GNW and IIT, and test them empirically in single unit recordings in non-human primates. Of note, we must emphasize that the predictions derived below, are according to IIT mathematics and are logical consequence to IIT and GNW literature, yet are not necessarily put forward explicitly by either IIT or GNW theorists. Further, these predictions bear on the nature of neural processing according to these theories, and are mute regarding “what it feels like” [24] or the “Hard Problem” [25] of consciousness. With these caveats in mind, first we formalize the role of multisensory neurons that integrate information from multiple sensory modalities (operationalized as being driven by multisensory stimulation than to unisensory stimulation, “AND” gates) vs. those that converge yet do not integrate (operationalized as responding to multiple sensory modalities but not being further driven by multisensory conditions, “XOR” gates = “OR” gates – “AND” gates; see [26] for an early characterization of multisensory neurons as Boolean gates). Interestingly, IIT mathematics suggests that a simple 3-node network (e.g., unisensory audio node, unisensory tactile node, and multisensory audio-tactile node) merging on an “AND” gate bears a greater degree of integrated information than one converging on an “XOR” gate (Φ = 0.78 vs. Φ = 0.25, respectively; see *Supplementary Information* online, Figure S1 & S2). Thus, according to this mathematical observation, it can be argued that as organisms’ transition from consciousness to unconsciousness, neurons capable of integration should be those most readily impacted, hence providing us with the first testable prediction (Prediction #1). Second, again derived from IIT mathematics, it may be suggested that when organisms are conscious, neurons that integrate information should demonstrate neural properties that are present during consciousness (see below) to a greater degree than do neurons that simply convergent information (Prediction #2). Finally, according to GNW, neural ignition, indexed as single trial co-activation, should be more readily apparent in conscious than unconscious states (Prediction #3).

To probe these predictions we simultaneously record single units from the primary somatosensory cortex (S1) and ventral pre-motor cortex (vPM) of non-human primates as they were presented with audio, tactile, or audio-tactile stimuli. The monkeys were trained to report the presence of a stimulus (regardless of sensory modality) via button press in order to determine their trial-to-trial alertness during propofol-induced loss of consciousness (see [27]). We first characterize both the central (e.g., mean) and dispersion (e.g., variance) tendencies of multisensory responses in S1 and vPM neurons under normal wakefulness, and based on these responses, divide neurons into integrative or convergent categories. We then describe the impact of anesthesia on neuronal ascription to these categories as the animals lose consciousness (testing prediction #1) and the degree to which they exhibit two neurophysiological indices that vary with consciousness - complexity [28] and noise correlations [29] (testing prediction #2). Finally, the fact that recordings were performed simultaneously within a known microcircuit (S1 & vPM; [30]) allowed us to examine single trial neural ignition as a function of consciousness (testing prediction #3).

## Results

### Characterizing Multisensory Neurons in S1 and vPM

The data, drawn as a subset of a previously published dataset [27], comprise neural responses recorded from 293 single units in S1 (228 from Monkey E and 65 from Monkey H) and 140 single units in vPM (87 from Monkey E and 53 from Monkey H) recorded across 26 sessions (16 in Monkey E and 10 in Monkey H). Responses were recorded as the animals were presented with either combined audiotactile (AT), tactile only (T), auditory only (A), or no (N) stimuli, and monkeys performed a detection task in which they were asked to press and hold a button for 3 seconds after sensory stimulation. Animals were progressively anesthetized with Propofol, at a rate determined to induce a loss of consciousness within approximately 10 minutes (see [27], and *Methods*). Thus, given the pattern of behavioral responses, trials can be divided into bins according to whether the animal was in an aware and unaware state (see Methods).

### Firing Rates in S1 and vPM neurons

Regarding the basic characterization of multisensory neurons in S1 and vPM, firing rates demonstrate 1) a reliable response to stimulus onset (Figure 1, 1^st^ and 3^rd^ row; colored horizontal bars indicate evoked response vs. baseline at p<0.01), and 2) a reduction in activity when monkeys were rendered unconscious, both with regard to spontaneous (i.e., baseline,) and evoked activity (Figure 1, 1^st^ and 3^rd^ row; shaded area represents the difference between evoked activity when animals were conscious and not at p<0.01). Further, as expected given the known role of vPM in auditory processing [31], neurons in vPM, but not S1, generally responded to auditory stimulation (interaction at p<0.01 between 60-210 ms post-stimuli onset).

**Figure 1.**
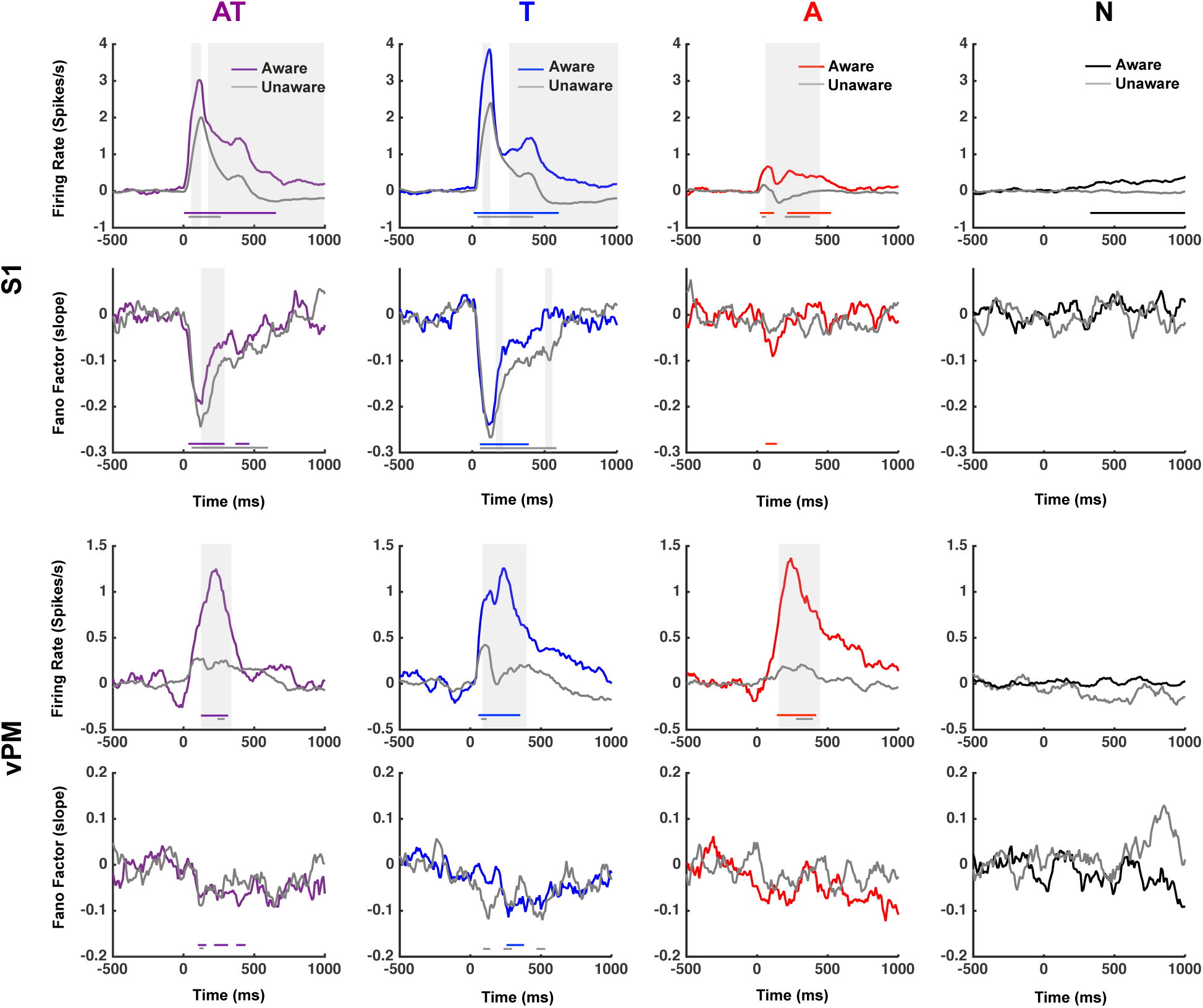
Time-resolved firing rates and Fano Factors in S1 and vPM as a function of state of consciousness. Presentation of audiotactile (AT; purple) and tactile (T; blue) stimuli evoked a reliable response in S1, while additionally the presentation of auditory (A; red) stimuli evoked a reliable response in vPM but not S1. Catch trials (N; black) did not evoked an increase in neural responses vis-à-vis baseline firing rate (0 on y-axis). Fano Factors were generally larger under states of unawareness than awareness (not depicted) and interestingly stimuli onset (0 on x-axis) quenched variability in S1 (particularly onset of AT and T stimuli) but less so (and not differently between states of consciousness) in vPM. The time-periods demonstrating a significant difference in evoked activity/FF as a function of state of consciousness (aware = colored, unaware = gray) are shaded in gray, while periods demonstrating a significant response vis-à-vis baseline are indicated by horizontal lines in each panel.

### Fano Factors in S1 and vPM neurons

Fano Factors (FF) were calculated to assess inter-trial response variability as a function of brain area, stimulation type, and state of consciousness. It has been previously reported that FFs are larger in an unconscious states (∼2.2) than in conscious states [29]. Our results support a similar pattern, with FFs values being on average 1.45 under unconscious conditions and 1.16 under conscious conditions. This pattern fits nicely with the notion that conscious neural representations are more reproducible than unconscious ones [32]. The results also show an interesting pattern of variability changes as a function of stimulus onset. Whereas prior work has shown a reduction in variability upon stimulus onset [33], the current results illustrate larger reductions in FF upon stimulus presentation in unconscious as opposed to conscious states (see *SI* for detail). This observation illustrates that firing rates and FFs are not directly yoked to one another (see [29], for a similar argument), and thus emphasizes the need to examine both central and dispersion tendencies in the response profiles of these neural populations.

### Multisensory Characteristics of S1 and vPM neurons

Following the observation that neural responses to stimulus presentation were most robust during the 500 ms immediately following stimulus onset (above, see *SI* for detail), we performed a spike count during this interval to characterize the multisensory properties of neurons recorded (see [34] for a similar approach). The multisensory integrative responses of these neurons were quantified in two ways: relative to the sum of the unisensory responses (i.e., relative to the additive prediction; Supra-Additivity Index), and 2) relative to the most effective unisensory response (i.e., quantifying the magnitude of the enhancement or depression of response; Enhancement Index [34, 35]. Here we report both indices for completeness, but operationally define integrative neurons as those with an enhancement index >1 (as both previous reports and the current dataset indicate that supra-additivity is rare in cortex (e.g., [36, 37]).

From the 293 single units recorded in S1, when the animals were aware 2 had a supra-additive index above 1 (supra-additivity index = 1.23 and 1.01, former depicted in Figure 2a), while another 2 (different neurons) had supra-additive responses when the monkeys were rendered unconscious (supra-additivity index=2.01 and 1.01). Thus, supra-additivity is seemingly rare in S1. On the other hand, 100 neurons had enhancement indices above 1 when the animals were conscious (see Figure 2b, for example), a number that was reduced to 55 when the animals were rendered unconscious (25 of which indicating an Enhancement Index greater than 1 in both aware and unaware states). Regarding vPM neurons, of the 140 neurons recorded, when the animals were aware 2 had a supra-additive index above 1 (Supra-Additivity Index=1.03 and 1.04), while only a single neuron was supra-additive when the animals were unconscious (Supra-Additivity Index=1.16, distinct neurons in aware and unaware cases). Twenty-nine vPM neurons had enhancement indices above 1 when animals were aware, a number that remained stable at 29 when monkeys were unconscious (2 out of the 29 neurons were the same in aware and unaware states). Hence, multisensory supra-additivity appears equally infrequent in S1 as in vPM, and interestingly there are seemingly more neurons demonstrating multisensory enhancement in S1 than vPM when animals were conscious (S1=34%, vPM=20%) yet approximately equal proportions when the animals are unconscious (S1=18%, vPM=20%).

**Figure 2.**
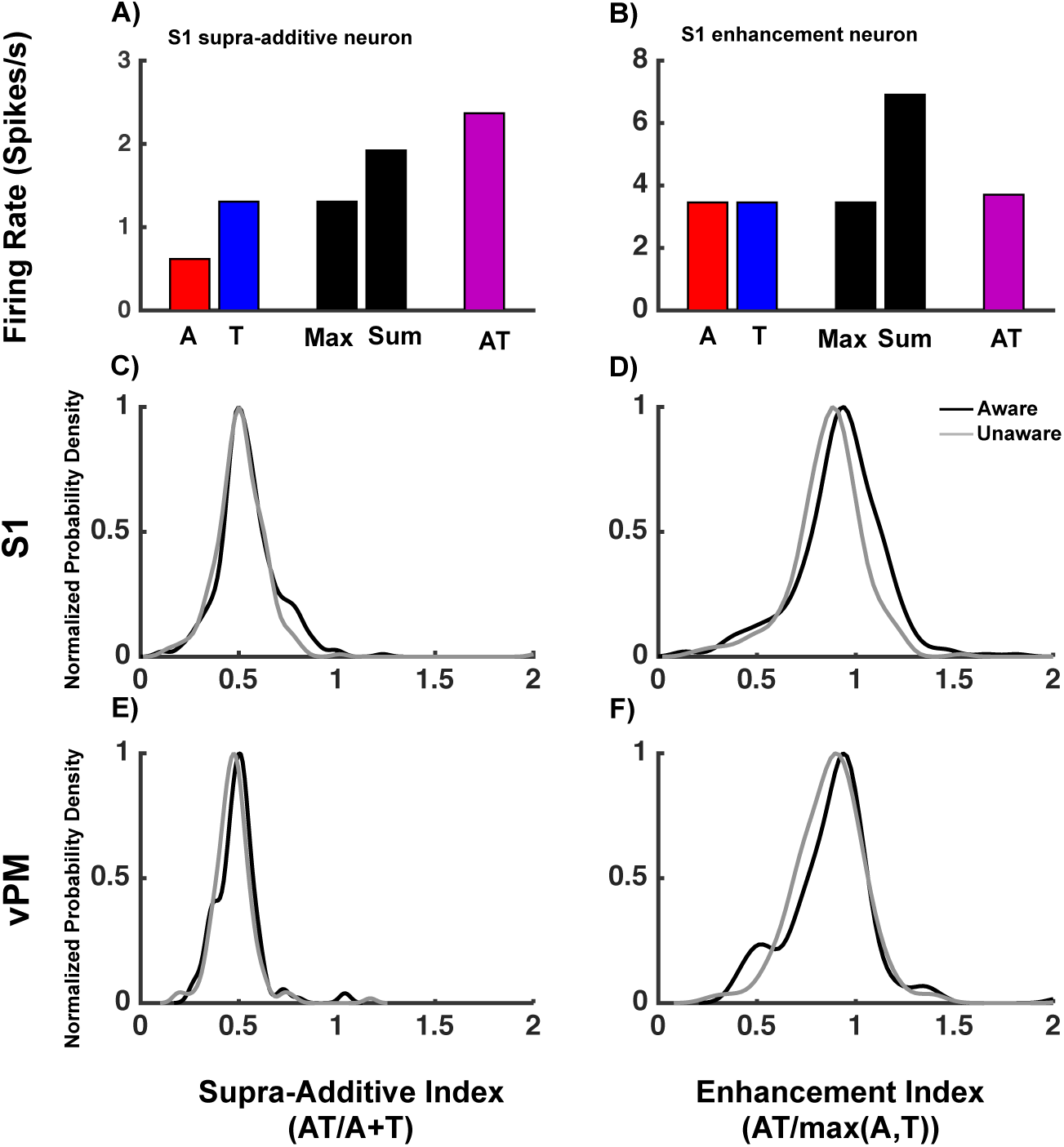
Characterizing multisensory neurons. A neuron whose multisensory response is greater than the sum of unisensory responses is said to be supra-additive (see A), while if it’s greater than the greatest unisensory response it’s considered to demonstrate multisensory enhancement (see B). A and B illustrate firing rates above a spontaneous rate (baseline-correction from −500 to 0ms; y-axis = 0). The distribution of supra-additive indices (left column) and enhancement indices (see Methods for detail) were normally distributed both in S1 (C and D) and vPM (E and F), regardless of whether the animals were aware (black) or unaware (gray).

The distributions of supra-additive and enhancement indices are well described by a Gaussian distribution, both when monkeys were conscious (chi-square goodness-of-fit test, p=0.28 and p=0.24, respectively) and unaware (p=0.70, and p=0.90, respectively; see [36] for a similar observation; Figure 2c-f). Given this, we can readily estimate the mean supra-additivity and enhancement indices associated with each neural population and consciousness state (see for example [38], for an indication that frequency of multisensory neurons and degree to which integration occurs may be dissociated). A 2 (recording area; S1 vs. vPM) × 2 (consciousness state; aware vs. unaware) independent samples ANOVA on supra-additive indices (Figure 2c,e) revealed main effects both of consciousness state (p=0.015) and recording area (p<0.01), where supra-additivity indices were larger under aware (M=0.52, S.E.M=0.006) than unaware (M=0.50, S.E.M=0.007) states, and larger in S1 (M=0.52, S.E.M=0.006) when compared with vPM (M=0.48, S.E.M=0.006). There was no interaction between these variables (p=0.47). On the other hand, a similar analysis with regard to enhancement indices suggested no distinction between consciousness states (p=0.11), no main effect of recording areas (p=0.30), and no interaction between these variables (p=0.09). These results highlight that the frequency and magnitude of multisensory integration may be dissociated (e.g., [38] and that supra-additive – where both unisensory responses are considered – and enhancement indices – where only the maximal unisensory response is compared to the multisensory response – may provide very different views into integrative capacity. Further, the findings indicate a highly heterogeneous population. Taking the example of the enhancement indexes in S1 (from which a representative multisensory integrative pool may be drawn, N=100, vs. N=29 in vPM) this metric indicates no overall change in the amount of integration at a population-level and across states of consciousness, yet examination of the classification of particular neurons reveals dramatic differences; shifting from 100 to 55 neurons in S1, only 25 of which were classified as integrating information both in aware and unaware states.

Fortunately within the current context aimed at examining theories of consciousness (e.g., IIT) we can leverage this variability to examine the outcome of neurons labeled as integrative or as convergent when animals are rendered unconscious. Figure 3 depicts the non-mutually exclusive compartmentalization of integrative and convergent neurons when monkeys were conscious. In this categorization scheme, neurons with an enhancement index greater than 1 were considered to integrate information. In the conscious state, 43% of neurons in S1 respond to both audio and tactile stimuli, and thus can be categorized as convergent (figure 3 top left). For vPM, this value is 44% (figure 3 bottom left). When the categorization is done based on integrative criteria, 52% of neurons in S1 were found to integrate auditory and tactile information (i.e., respond to AT+(AT> max (A, T); figure 3 top right), while 33% of neurons in vPM were categorized as integrative (figure 3 bottom right).

**Figure 3.**
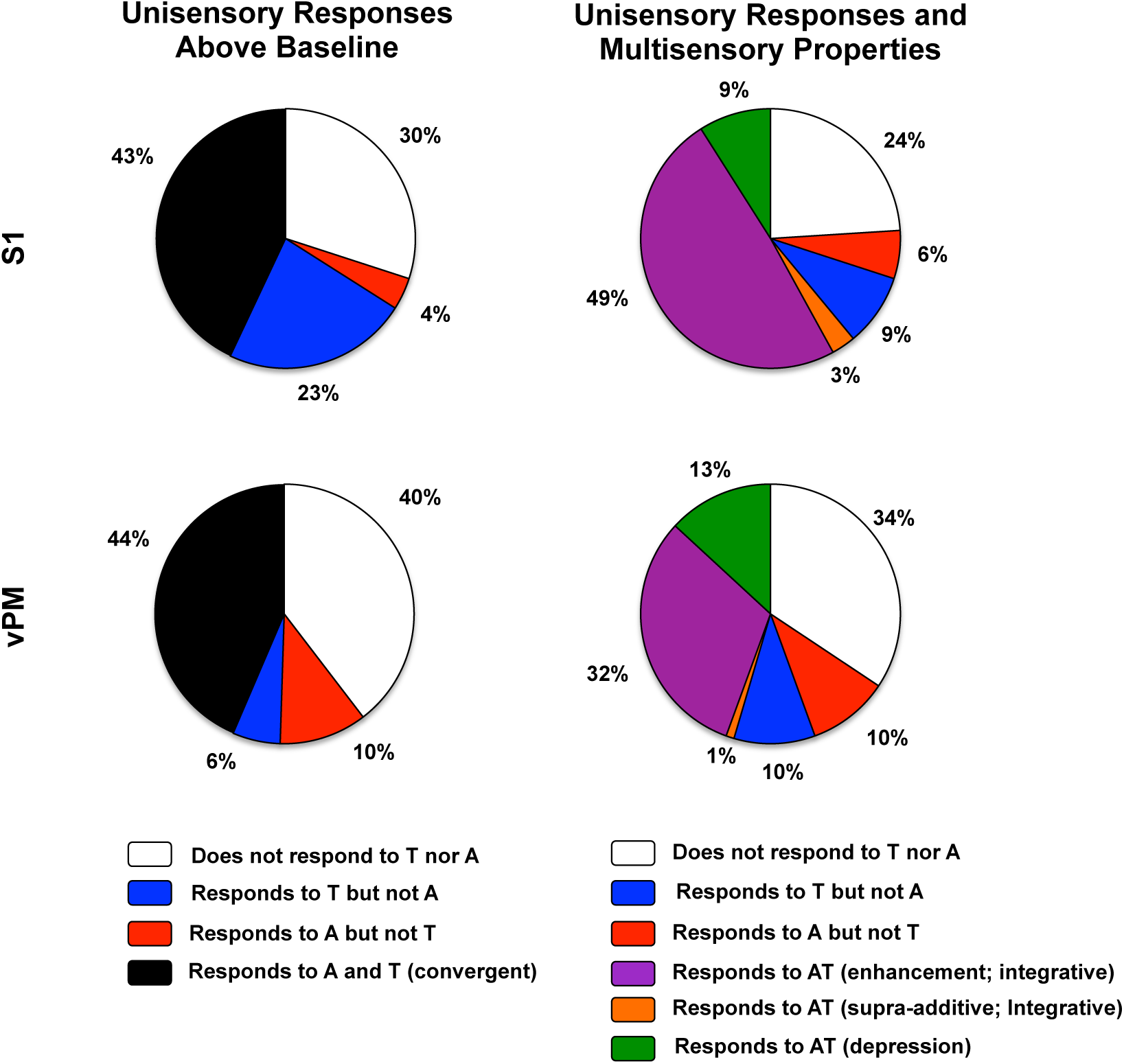
Non-mutually exclusive classification of neurons in S1 (top row) and vPM (bottom row) as convergent, integrative, unisensory, or non-responsive. Left column; Neurons whose convolved firing rate excited their spontaneous rate plus 2 standard deviations for at least 10ms between 0ms and 1000ms post-stimuli onset were responsive. If they responded to both tactile (T) and auditory (A) stimulation, they were considered convergent (black). On the other hand, if they responded solely to T or A stimulation, they were respectively labeled as tactile (blue) and auditory (red). Right column; Differently from the case of convergence, in order to characterize a neuron as integrative, their response profile to audio-tactile (AT) stimulation had to be examined. First, neurons were classified as responsive or not (white; as above). Next, if the neuron was responsive to AT stimulation (defined as above) we queried whether during some epoch between 0ms and 1000ms post-stimuli onset their firing rate to AT stimulation was greater than the sum of A and T firing rates (supra-additivity; orange) or the maximum of A and T firing rates (enhancement; purple). Lastly, if a neuron was responsive to AT stimulation but responded less to AT than to unisensory stimulation, the neuron was classified as demonstrating multisensory depression (green). Lastly, if they neuron did not respond to AT or A stimulation, but did to T, it was labeled as tactile (blue), while if a neuron did not respond to AT or T, but did to A, it was labeled as auditory (red).

Importantly, in order to examine how this categorization is changed when animals are rendered unconscious (Prediction #1) and to quantify the extent to which they exhibit properties of consciousness (Prediction #2), we created mutually exclusive groups. Neurons that failed to respond to the auditory and tactile stimulus combination more that to the individual stimuli were classified as strictly convergent (*convergent yet not integrative*, or “XOR” gates). Conversely, neurons that responded more vigorously to the multisensory combination were labeled as integrative neurons, and were operationally categorized as “AND” gates. This bifurcation of neurons into exclusive groups is important from a statistical perspective (in order not to create groups that are partially overlapping and overlapping to different extents across states of consciousness and recordings areas), and most importantly, from a theoretical perspective, creating “AND” and “XOR” neuronal pools. However, given the initial number of recorded neurons in S1 and vPM, this categorization scheme yielded a sufficient number of convergent (N=125) and integrative (N=64) neurons in S1, but not in vPM (convergent, N = 61; integrative, N = 8). Thus, for the analyses specifically probing the difference between convergent and integrative neurons in light of IIT (Predictions #1 and #2), analyses are restricted to S1.

### Testing Consciousness Theory in Multisensory Neurons; Information Integration Theory Prediction #1; Are integrative neurons most readily impacted by loss of consciousness?

A first neurophysiological prediction that may be derived from the IIT is that network structured around an integrative neuron should lead to a greater degree of consciousness than one structured around a convergent neuron (see *SI and Introduction*). Hence, as an organism is rendered unconscious, the prediction is that integrative neurons should be most impacted.

As illustrated in Figure 4a, while a significant portion of S1 neurons labeled as convergent when the monkey was conscious became responsive exclusively to touch (42.1%) following loss of consciousness, others were rendered non-responsive (24.1%) or transitioned to responding exclusively to auditory stimulation (2.5%). 31.0% of neurons remained responsive to both auditory and tactile stimulation following loss of consciousness. On the other hand, of S1 neurons labeled as integrative when the animal was conscious, nearly two-thirds (62.9%) remained integrative following the loss of consciousness. 18.6% of integrative neurons became exclusively responsive to tactile stimulation, 2.2% became exclusively responsive to auditory stimulation, and 16.3% became unresponsive. A Chi-squared test demonstrated that these proportions (62.9% remaining as integrative but only 31.0% remaining as convergent) were significantly different from one another (p=0.001). Thus, and in contrast to the prediction derived from IIT, convergent neurons were more impacted when monkeys became unaware. It must be noted that this occurred despite the fact that arguably the requirements for being classified as “integrative” (i.e., responding to AT stimuli beyond their spontaneous and responding to AT stimuli beyond the maximal unisensory response) was more stringent than the bar required for a neuron to be classified as “convergent” (i.e., responding to A and T stimuli beyond their spontaneous firing rate).

**Figure 4.**
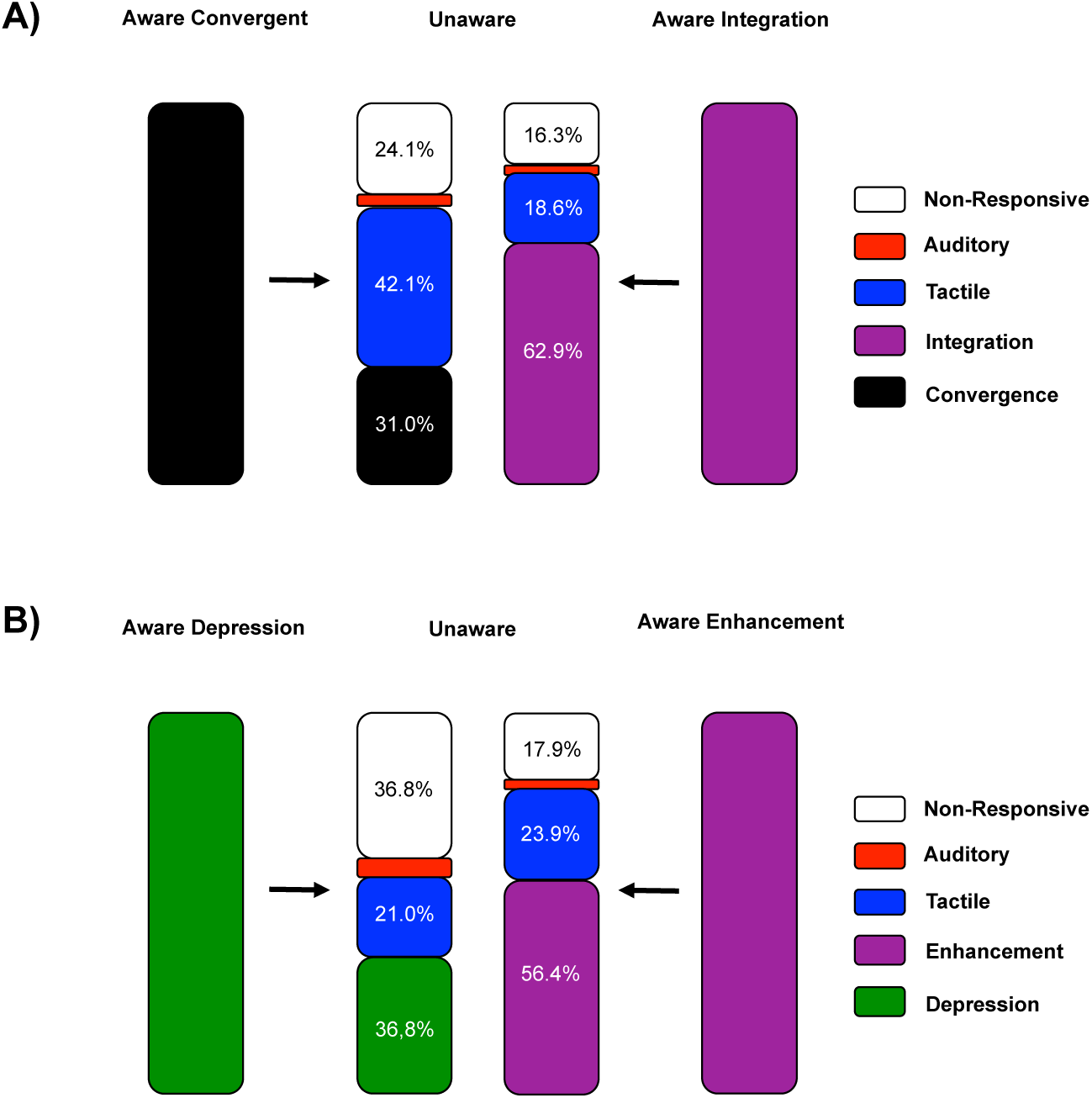
Transitions of S1 convergent and integrative neurons into distinct categories as monkeys are anesthetized. **A)** The largest proportion of convergent neurons when monkeys were aware (black, leftmost) became responsive solely to tactile stimulation (blue) when monkeys were rendered unconscious (second column), while 31.0% remained as convergent (black, second column). On the other hand, the majority of integrative neurons when monkeys were aware (rightmost column) remained as integrative (purple, 3^rd^ column). **B)** Similar to the contrast between convergent and integrative neurons, when contrasting neurons exhibiting multisensory depression (i.e., responds to AT but to a lesser extend than to unisensory stimulation) and enhancement (i.e., responds to AT and to a greater degree than to unisensory stimulation), results suggests that the larger the multisensory gain, the more neurons remain as integrative (vs. not) when rendered unconscious.

We further examined whether these anesthesia-induced changes in neuronal responsiveness scaled with the degree to which neurons may be considered to be integrative. While supra-additivity or multisensory enhancement are considered to be the hallmarks of multisensory integration [27], many multisensory neurons respond less vigorously to multisensory stimulation than their maximal response to unisensory stimulation (i.e., multisensory depression; [35]. Nonetheless, these neurons are still considered to play an important role in multisensory integration [39]. As illustrated in Figure 4b, while 56.4% of neurons exhibiting multisensory enhancement during consciousness had this enhancement preserved when the animal was rendered unconsciousness, only 36.8% of neurons that were categorized as exhibiting multisensory depression remained in that category upon the transition to unawareness. These proportions were significantly different from what is expected under the null distribution (p=0.04). In sum, not only are integrative neurons not most readily impacted during the loss of consciousness, but the more a neuron is driven by paired stimulation toward response enhancement, the more likely it is to retains this enhancement during unconsciousness.

### Prediction #2; Do integrative neurons most readily demonstrate neural properties associated with consciousness?

In addition to probing the fate of convergent and integrative neurons as the animals were rendered unconscious, we also probed the degree to which these neurons exhibit neurophysiological properties associated with conscious states. The empirical measure most commonly associated with the IIT is the perturbation complexity index (PCI; [6]) and a component of this index, Lempel-Ziv complexity (LZ; [28]). In short, PCI is calculated by perturbing the cortex via transcranial magnetic stimulation (TMS) in an attempt to engage a distributed brain network, and subsequently compressing the spatiotemporal patterns of neural activity generated by the perturbation (using LZ) to measure the complexity of the response. In theory, the more distributed and recurrent the network, the larger should be the spatiotemporal complexity evoked by the perturbation. This empirical measure was directly derived from the IIT [11] and has been shown to successfully differentiate between distinct levels of consciousness [6, 40]. In more simple applications, LZ has also been applied to resting state [41, 42] and stimulus evoked [43, 44] neural activity (most commonly in scalp EEG datasets), and similar to PCI, has been shown capable of differentiating between levels of consciousness [41, 42]. Here we first characterize time-resolved LZ complexity in spike trains as a function of consciousness state and modality of stimulation. The analysis is performed both on baseline-corrected values (in order to compare the changes evoked by sensory stimulation) and on non-baseline-corrected values (in order to more generally examine the relationship between LZ complexity in spike trains and level of consciousness). After establishing predictions based on the state of consciousness in the neural population as a whole, we then bifurcate S1 neurons into convergent or integrative pools, and examine which cohort most faithfully exhibits LZ complexity values that tracks the animal’s consciousness state.

As illustrated in Figure 5A, overall LZ complexity was greater (across the entire epoch, see *SI*) when monkeys were unaware (Figure 5, 1^st^ and 3^rd^ rows respectively for non-baseline corrected LZ in S1 and vPM) as opposed to aware. In addition, stimulus evoked suppression of complexity (Figure 5, 2^nd^ and 4^th^ rows) was more sustained under conscious than unconscious conditions, particularly in S1 (Figure 5A, shaded areas are significantly different between consciousness states and horizontal colored lines in baseline corrected panels show interval of evoked reduction in complexity, see *SI* for detail). These general properties of LZ complexity were next indexed in convergent and integrative neurons. As depicted in Figure 5B, a 2 (consciousness state; aware vs. unaware) × 2 (neuron type; convergent vs. integrative) ANOVA on non-corrected values demonstrated a main effect of awareness (aware; M=0.80, S.E.M=0.001; unaware; M=0.87, S.E.M=0.002; p<0.01), yet no main effect of neuron type (all p>0.11). Most interestingly, however, there was a significant interaction between these variables (p<0.01), as convergent neurons (M=0.79, S.E.M.=0.002) had marginally lower LZ complexity than integrative neurons (M=0.81, S.E.M.=0.002) when monkeys were aware (p=0.052), yet this pattern reversed when monkeys loss consciousness (integrative; M=0.86, S.E.M.=0.002; convergent; M=0.88, S.E.M=0.001, p=0.045). Thus, when quantified using uncorrected LZ values, convergent neurons tracked the state of consciousness – i.e., they exemplified the LZ behavior expected from a given state of consciousness (Figure 5A) – better than did integrative neurons. A similar analysis corrected for different baselines indicated a main effect of consciousness state (p<0.01 between 50ms and 700ms post-stimuli onset), but failed to indicate a difference between neuron types (all p>0.02), or an interaction between these variables (all p>0.09). Hence, while the overall level of LZ complexity appeared to differentiate between convergent and integrative neurons, the duration and/or magnitude of the change in LZ complexity during evoked responses did not.

**Figure 5.**
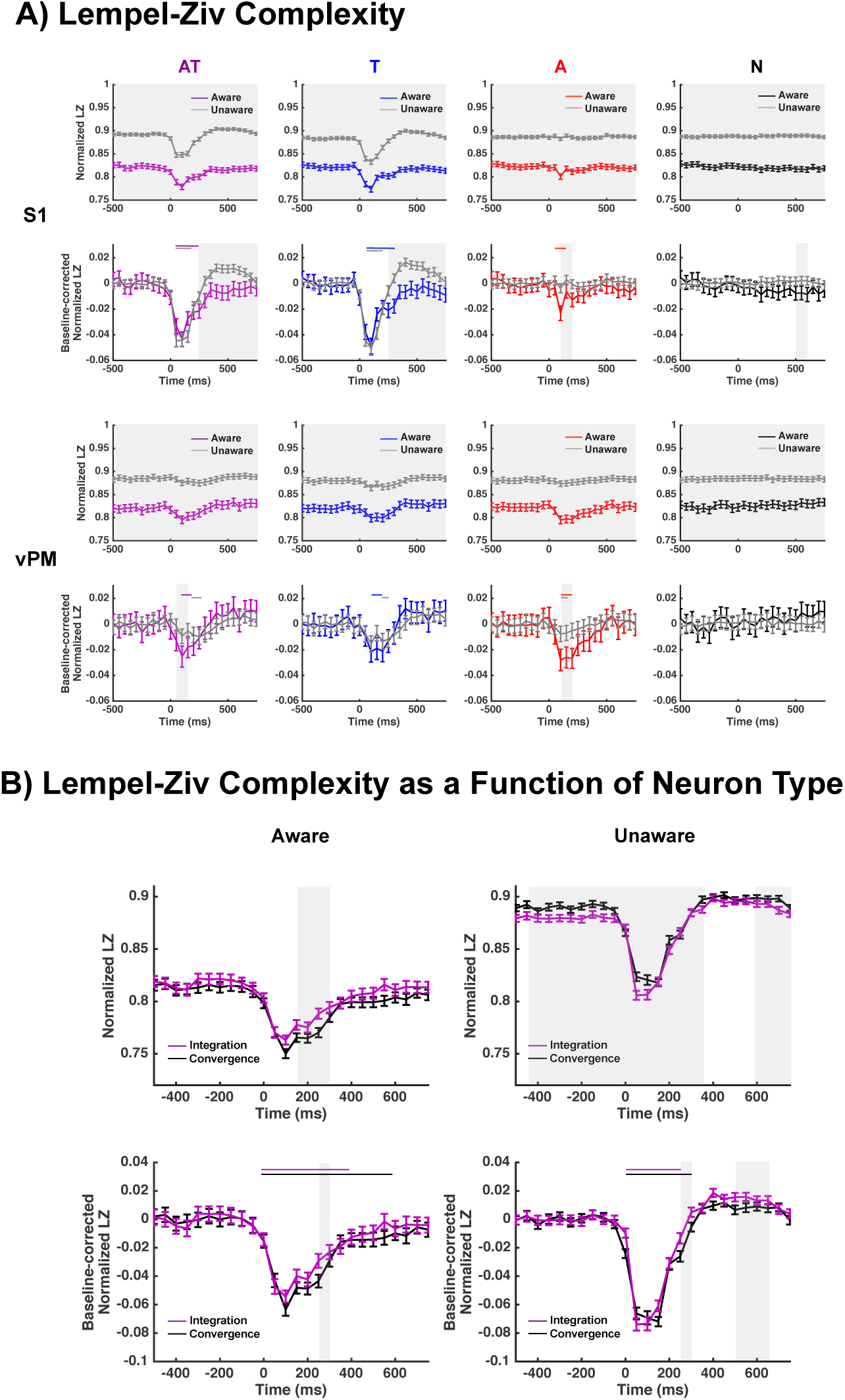
A; Time-resolved evoked Lempel-Ziv Complexity in spiking activity in S1 (top 2 rows) and vPM (bottom 2 rows) neurons as a function of consciousness state (aware = colored; unaware = gray) and sensory stimulation (AT = purple, T = blue; A = red; none = back). Most strikingly as illustrated when time-courses were not corrected for baseline (1^st^ and 3^rd^ rows) results suggest an increase in complexity (y-axis) when monkeys were rendered unconscious. Further, as better exemplified when correcting for baseline (2^nd^ and 4^th^ rows), the evoked complexity (negative deflection) is seemingly more sustained when aware than unaware. **B; LZ in spiking activity in S1 neurons as a function of consciousness state (aware = 1**^**st**^ **column; unaware = 2**^**nd**^ **column) and whether the neuron was determine to converge (black) or integrate (purple) sensory information when aware.** Results suggest that normalized LZ (top row, y-axis) is higher for integrative than convergent neurons when monkeys are aware (left column) yet this pattern reverses when monkeys are rendered unconscious (right column). Similarly, the evoked nature of LZ complexity due to AT stimulation (bottom row) was similarly more sustained for convergent than integrative neurons (particularly when aware; left column), however there was no significant interaction between consciousness state and neuron type when normalized LZ complexity was corrected for baseline.

These complexity results, just like the observed shifts in the distributions of convergent and integrative neurons following loss of consciousness, point in a direction counter to IIT – in that they suggest that convergent, as opposed to integrative, neurons more faithfully exhibit properties of consciousness. However, an important caveat is that there is relatively little empirical work quantifying LZ complexity in spike trains [45, 46]. Hence, it may be useful to apply a similar logic – contrasting convergent and integrative neurons as a function of consciousness – while utilizing a better-characterized neurophysiological measure within the context of consciousness studies. Thus, we next examined noise correlations.

Noise correlations – the degree to which the response of a pair of simultaneously recorded neurons co-vary after accounting for the signal – were originally considered to originate from shared sensory noise arising in afferent sensory pathways [47]. However, more recent studies suggest that correlated noise may also reflect meaningful top-down signals generated internally within the central nervous system [48]. Most interestingly for the current work, noise correlations have been shown to be strongly dependent upon state of awareness. For example, one study has demonstrated a six-fold increase in these correlations under an opioid anesthetic when compared to wakefulness (unaware = 0.05; aware = 0.008; [29]). Accordingly, we examined noise correlations as a function of recording area (S1 and vPM), type of sensory stimulation (AT, T, A, and N), and consciousness state (aware and unaware). In addition to their relevance to the predictive framework laid out in the *Introduction* (specifically prediction #2), our assessment of noise correlations in our data set has added importance as it represents the first measure of the impact of propofol on single unit noise correlations in non-human primates.

As illustrated in Figure 6A, noise correlations demonstrated a striking increase from aware (M = 0.02, S.E.M = 0.001) to unaware (M = 0.11, S.E.M = 0.002) states (F=742.76, p<0.001). This effect was independent of recording area (p = 0.86) and stimulation type (p = 0.33), nor was there an interaction between variables in driving the degree to which noise correlated across single units (all p>0.11). Thus, the current dataset (utilizing Propofol) is in general agreement with the opioid-derived observation [29] in that under anesthesia noise correlations increase by approximately six-fold.

**Figure 6.**
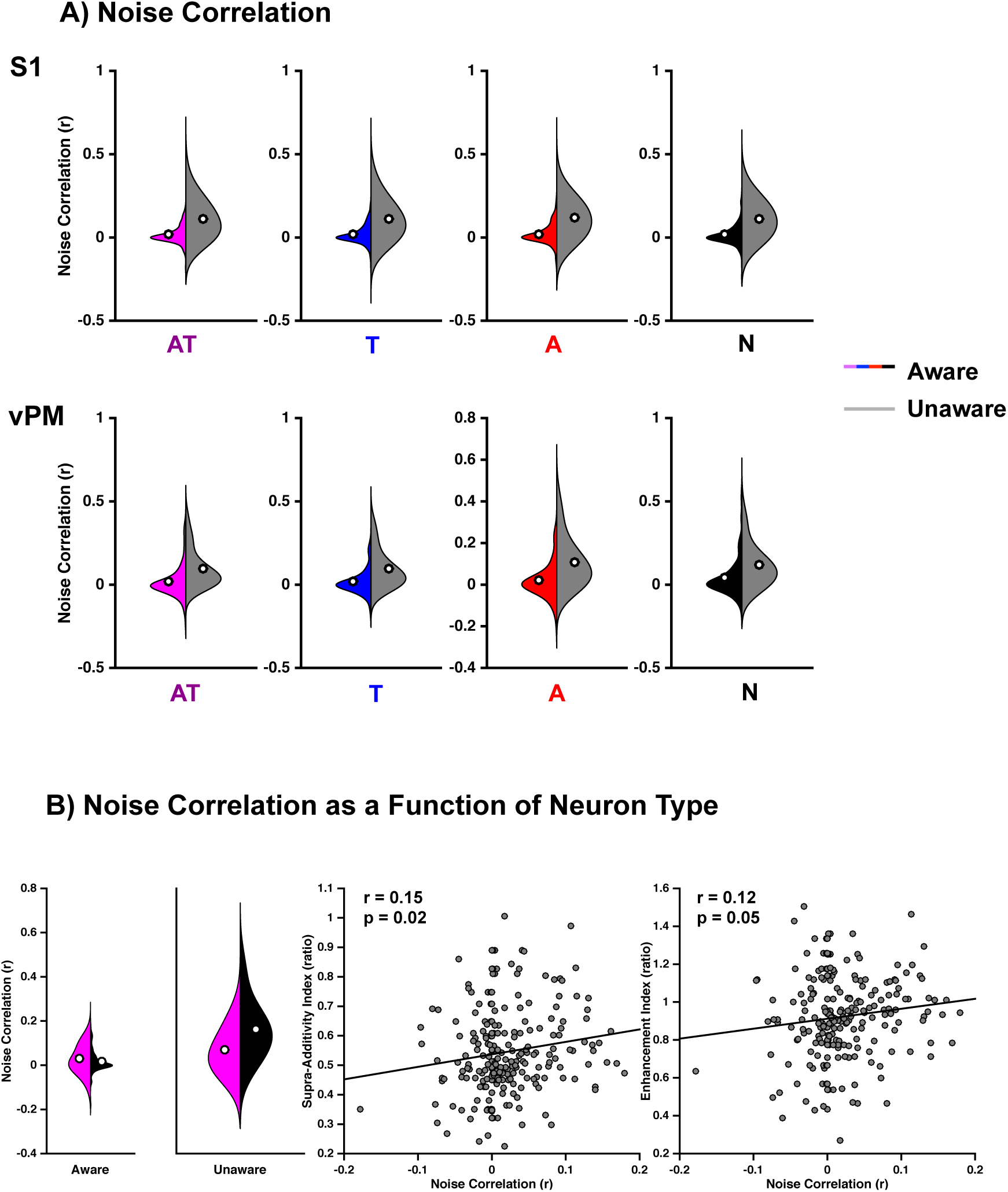
A; Noise correlations in S1 (top) and vPM (bottom) as a function of consciousness state and sensory stimulation. Violin plots colored (purple = AT, blue = T, red = A, black = N) represent conscious states, while their gray counterparts illustrate noise correlations when the monkeys were rendered unconscious. White dots emphasize the mean. Overall, across all sensory modalities, noise correlations are 6-fold greater under unconscious than conscious states. **B; Noise correlations in integrative and convergent S1 neurons.** When monkeys are aware (leftmost panel) integrative neurons (purple) exhibit a higher degree of noise correlations than neuron that integrate (black), while the contrary is true when monkeys were rendered unaware (2^nd^ column). Further, when monkeys were aware, the more a neuron exhibited noise correlations (3^rd^ and 4^th^ panel, x-axis) the greater it’s supra-additive (3^rd^ panel, y-axis) and enhancement (4^th^ panel, y-axis) indices. White dots represent the mean of each distribution.

When restricting noise correlation analysis to the integrative and convergent neurons, we observed significant main effects of consciousness state (F=91.56, p<0.001) and neuron type (F=19.59, p<0.001), as well as an interaction between these variables (F=29.63, p<0.001). The interaction seems to be driven by the fact that when monkeys were unconscious convergent neurons (M=0.16, S.E.M=0.02) showed a greater degree of noise correlations than integrative neurons (M=0.068, S.E.M=0.008; p<0.001; see Figure 6B), and this difference disappeared during consciousness (Convergent: M = 0.018, S.E.M = 0.004; Integrative: M=0.028, S.E.M=0.008; p = 0.067), significant (p=0.067). Thus, while noise correlations were lower during conscious rather than unconscious states (in the population as a whole), this pattern is most readily apparent in convergent rather than integrative neurons.

Overall, the results suggest that consciousness is marked by a reduced degree of noise correlations, and that integrative neurons poorly track level of consciousness as indexed by this measure. In fact, these findings suggest a potential negative relationship between the degree of noise correlation that is typically associated with consciousness on one hand and with integration on the other. Namely, neurons that demonstrate the greatest degree of integration are those that show the greatest degree of noise correlation (in the conscious state). In support of this hypothesis, and as illustrated in Figure 6B (middle and right panel), both the supra-additivity (r=0.15, p=0.02) and enhancement (r=0.12, p=0.05) indices were positively correlated with the degree to which a neuron exhibited noise correlations.

### Testing Consciousness Theory in Multisensory Circuits; Global Neuronal Workspace (Prediction #3)

Beyond the characterization of single cell properties (i.e., firing rate, Fano Factor, complexity), it is possible to leverage the fact that neurons in both S1 and vPM – a well-studied microcircuit [30] – were concurrently recorded to test another prominent theory of consciousness; the GNW theory [8]. This theory states that sensory stimuli will elicit a conscious percept when the neural activity associated with the stimuli propagates through a broad fronto-parietal network following neural ignition. Adapting this theoretical framework to the current experimental design, GNW predicts a higher likelihood of near concurrent neuronal firing in S1 and vPM (at the single trial level) when animals are in a conscious state (and thus capable of conscious content) than when they are unconscious (Prediction #3).

To test this prediction we define a response threshold as exceeding spontaneous firing by two standard deviations (see *Methods*), and then calculate the percentage of trials that result in significant firing in S1, vPM, or both S1 and vPM, as a function of consciousness and sensory stimulation type. This approach yields relatively small percentages of trials catalogued as “active”, which is to be expected given Poisson firing (i.e., the fact that on most trials the firing rates of most neurons change modestly, with relatively few neurons driving global population changes [33]), the high threshold set for labeling a trial as “active”, and the requirement for near concurrent firing. This approach, in other words, is statistically conservative. As highlighted in Figure 7 (leftmost panel), results revealed that when animals were conscious, during combined AT stimulation, both S1 and vPM were concurrently active on 1.17% of trials (labeled “Concurrent Activation”). This number is reduced to 0.96% of trials during T stimulation, to 0.67% of trials during A stimulation, and to 0.28% of catch trials (main effect of stimulation type during awareness; Friedman Test, χ^2^ = 135, p < 0.001). The percentage of trials in which sensory stimulation resulted in the co-activation of S1 and vPM was significantly smaller when animals were rendered unconscious (main effect of consciousness state, Wilcoxon Test, Z=1135, p<0.001) and did not differ across stimulation types (Friedman Test during unawareness; χ^2^=14.32, p=0.64; stimulation type by consciousness state interaction, Friedman Test of the difference between conscious vs. unconscious as a function of sensory stimulation type, χ^2^ =204.78, p<0.001). A similar pattern of results emerged when examining the number of trials that resulted in the independent activation of S1 and vPM (see *SI* for detail). Thus, in S1, 13.2% of AT trials resulted in significant firing when monkeys were conscious, a number that was reduced to 10.5% in T trials (Wilcoxon, p=1.61e-19), and further reduced to 6.5% in A and 6.1% in N trials (T vs. A, p=1.28e-8; A vs. N, p=0.68). In vPM, interestingly, the main effect of trial type in the conscious condition (see SI) resulted from AT, T, and A all being different from N trials (all p<5.0e-20), as well as from vPM firing being most likely due to A stimulation (M=8.4%) than to AT (M=7.4%) or T (M=7.5%) stimulation (all p<2.3e-5). That is, activation of vPM was more probable due to A stimulation than T or AT stimulation – a stipulation that was not true (in fact opposite) in S1 or when examining co-activation of S1 and vPM. This finding pinpoints that auditory information must arrive to vPM via a route that is not the same as how tactile information arrives in vPM (e.g., via S1), a finding that makes a great deal of sense since vPM is known to be part of the auditory “what” or ventral pathway [49]. Lastly, on the vast majority of trials sensory stimulation did not result in activity in either S1 or vPM, a finding that is most prominent in unconscious (M=91.0%) than conscious states (M=81.3%, Z=37949, p < 0.001).

**Figure 7.**
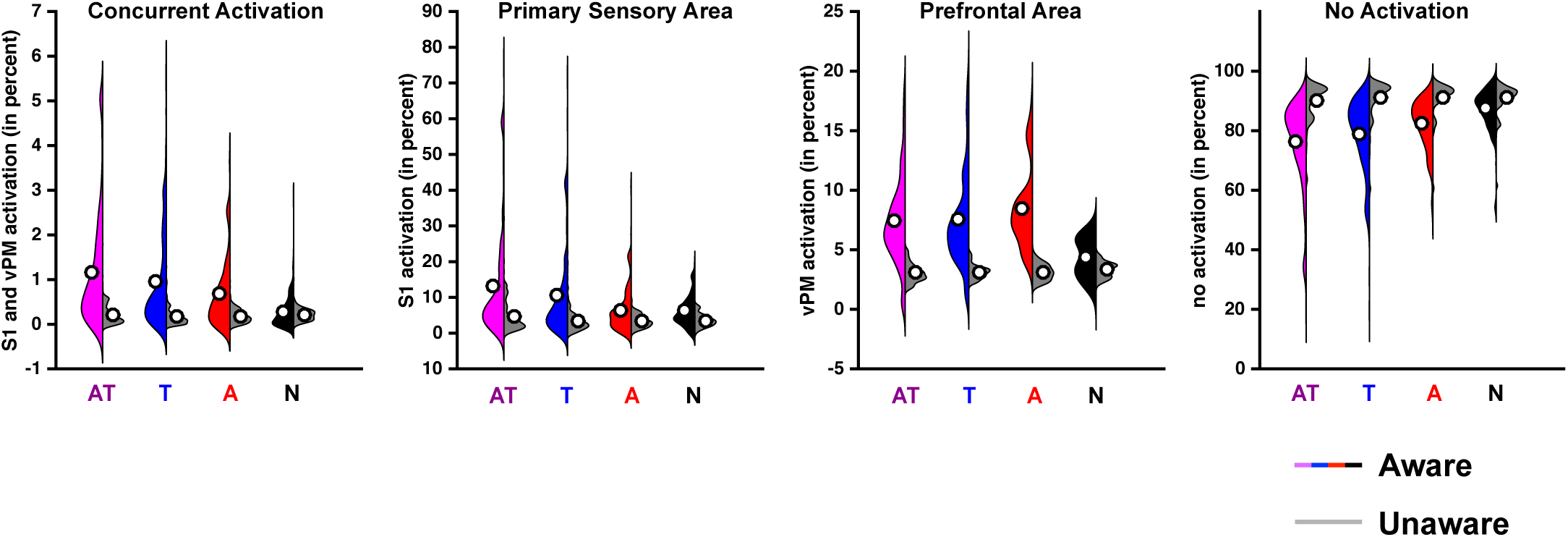
Percentage of trials that result in significant activation of S1, vPM, both or neither area, as a function of consciousness state and stimulation type. Concurrent activation is defined as the simultaneous activation of S1 and vPM (leftmost panel). This phenomenon occurs to a greater degree when animals were conscious than unconscious, during AT (purple), T (blue), or A (red) stimulation, but not during catch trials (no stimulation). 2^nd^ and 3^rd^ panel respectively demonstrate the number of trials that result in significant activation of S1 and vPM. Interestingly, while AT and T stimulation seemingly result in a greater percentage of trial demonstrating neural ignition and S1 activation than A stimulation, this is not the case for activation of vPM. Namely, auditory information seemingly reaches prefrontal areas via other routes. Lastly, rightmost panel illustrates the percentage of trials that do not result in significant activation; here the percentage is greater in unconscious than conscious trials, regardless of type of sensory stimulation. White dots represent the mean of each distribution.

Thus, the overall pattern of results incorporating all cells recorded illustrate that when monkeys were conscious and sensory stimuli were being presented a greater number of trials resulted in co-activation of both primary sensory and “associative” areas than when animals were unconscious. The finding is in line with the GNW theory, but may represent a trivial result given that a larger number of trials also show exclusive activation in S1 or vPM when the animals were conscious. Hence, for pure probabilistic reasons, co-activation of S1 and vPM would be more likely under conscious than unconscious conditions. To address this concern, in a second step of analysis, we multiplied the likelihood of observing activation in S1 by the likelihood of observing activation in vPM and contrasted this predicted value to that observed (for both conscious and unconscious conditions). As shown in Figure 8, results demonstrated that in both the aware (M=0.49%, one-sample t-test to zero, p=2.79e-22) and unaware (M=0.08%, one-sample t-test to zero, p = 2.95e-13) cases, co-activation of S1 and vPM was more likely than what would be predicted by simply multiplying probabilities (Figure 8, y = 0). More importantly, the degree to which co-activation exceeded this prediction was greater under conscious conditions than under unconscious conditions (t=6.2, p=6.41e-10).

**Figure 8.**
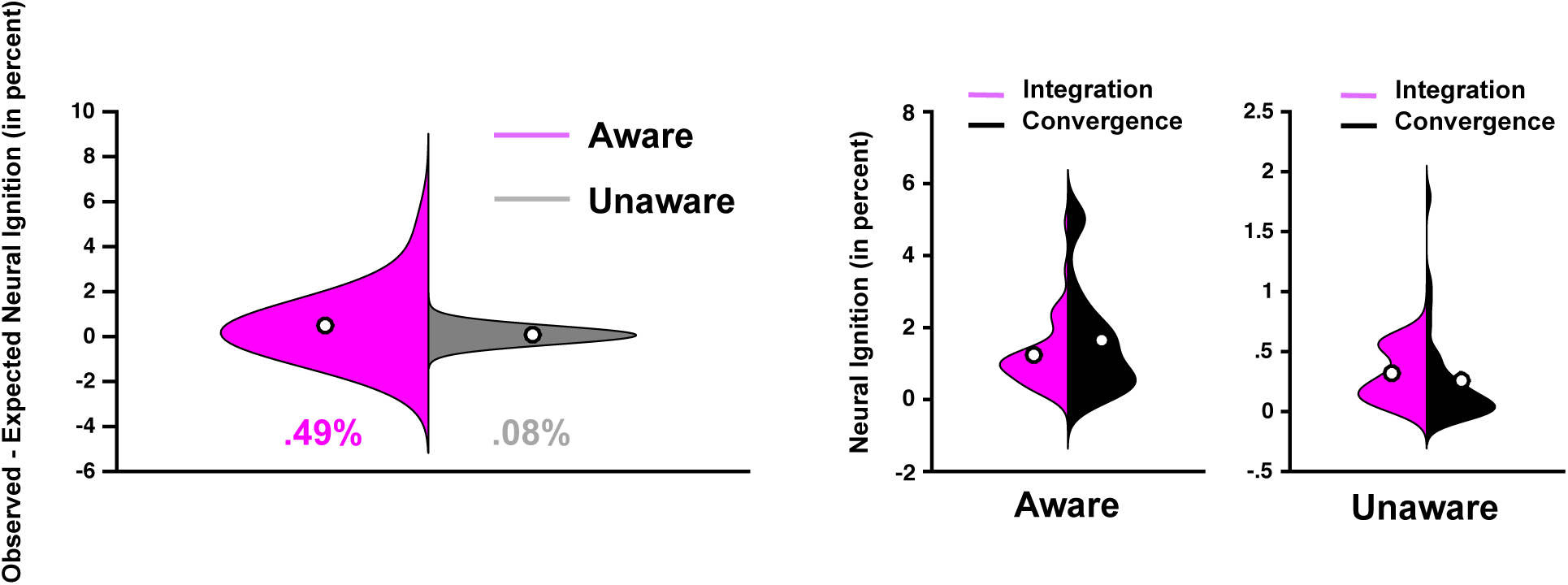
Neural ignition as a function of conscious state. **A)** Difference in the observed percentage of trials resulting in neural ignition due to AT stimulation from the percentage of trials that would be predicted based on S1 and vPM activation alone (y = 0), as a function of conscious state (aware = purple; unaware = gray). **B)** Neural ignition due to AT stimulation in integrative (purple) and convergent (black) neurons as a function of consciousness state (aware = left; unaware = right). White dots represent means of the distribution.

Lastly, as the previous results (Figures 5, 6; testing the IIT) had suggested that convergent neurons exhibited properties of consciousness to a greater degree than integrative ones, we sought to determine whether this was also true for these co-activation results. As illustrated in Figure 8 (center and right-most panels) co-activation was generally more common when monkeys were conscious (M=1.5%) than unconscious (M=0.3%, Mann-Whitney U, p<0.001). Further, these likelihoods interacted with neuron type. Co-activation was not distinct in convergent and integrative neurons when the animals were aware (convergent, M = 1.8%; integrative, M = 1.6%, p=0.37). In contrast, convergent neurons demonstrated less co-activation than integrative neurons when the animals were aware (convergent, M=0.26%; integrative, M=0.31%, p=0.004).

## Discussion

Detailing the neural mechanisms enabling wakefulness (i.e., consciousness-level) and conscious experiences (i.e., consciousness-level) is a central question within systems neuroscience [7]. As such, several theoretical frameworks have been put forth [7, 8, 11-17, 50]. Two of the most influential of these are Information Integration Theory (IIT; [11, 50]) and Global Neuronal Workspace (GNW; [7, 8]). Unfortunately, these theories have seldomly been tested neurophysiologically, and never within the same dataset. A series of concrete predictions can be generated from these theories, and in the current work we sought to generate such predictions and test them.

Starting from IIT mathematics, a strong prediction that can be made is that as an organism transitions from conscious to unconscious states, central integrative hubs (vs. convergent hubs) of neural networks should be most impacted. Indeed, IIT states that the greater the information possessed by a network above and beyond its constituent parts, the more conscious the system [11, 50]. Level of consciousness may be calculated and represented as the variable phi (Φ). We demonstrate that within a simple three-node network, if the central node is an integrative (“AND” gate) node as opposed to a convergent (“XOR” gate) node, the value of Φ triples (see *SI*). In evaluating the neuronal data, we categorized neurons as either convergent or integrative and examined which class was most impacted by propofol administration. The assumption here is that cross-modal neurons in S1 and vPM receive information regarding the different senses from upstream areas, S1 and vPM in turn being the central node composed of “AND” and “XOR” functionality. Of course, this is an over-simplified biological neural network, but one that permits testing predictions derived from the IIT from a neurophysiological perspective. Contrary to our IIT-derived predictions, convergent, as opposed to integrative, neurons were most impacted by the administration of anesthesia. To further test predictions derived from IIT, we reasoned that when organisms were conscious, integrative neurons should exhibit neurophysiological properties of consciousness to a greater extent than convergent neurons (i.e., supporting lower phi-values). The two measures chosen were Lempel-Ziv complexity and noise correlations, and we examined these as a function of stimulation type and conscious state. It is important to note that we do not aim to explain why neural complexity or noise correlations are altered by consciousness state, but simply to use these measures as “features of consciousness” and index how these properties are modulated in integrative vs. convergent neurons as a function of consciousness. Complexity was chosen as it is the measure most often used within the IIT framework (e.g., [41, 42] – though the relationship between Φ, Lempel-Ziv complexity, TMS-evoked complexity (PCI) and stimulus-evoked complexity is far from clear [51]). The study of noise correlations was chosen as this measure has a stronger tradition within neurophysiology, and prior studies [29] have shown substantial increases in noise correlations after administration of an opioid anesthetic. Here again, findings indicated that convergent neurons most closely tracked the animals’ consciousness state. Taken together, the findings of these neurophysiological analyses fail to provide strong empirical support for the IIT.

One of the novel findings of the current study is that under propofol – a GABAA potentiator [52] – noise correlations are approximately six-fold greater than during wakefulness. This finding is in line with previous single unit recordings under a different anesthetic (opioid; [29]) and concordant with a recent graph theory analysis of electrophysiological data showing that a change in local information processing efficiency – a measure that changes with noise correlations - could differentiate between distinct levels of responsiveness due to propofol administration [53].

The second theory tested was GNW [7, 8]. In GNW the core concept is that during wakefulness a conscious experience should result in neural ignition – the broadcasting of sensory evidence throughout the brain. Concordant with the basic tenets of GNW, our results suggest that the co-activation of primary sensory areas and higher-order areas on a single trial is more likely under conscious than unconscious conditions. Importantly, the occurrence of this co-activation exceeded the expected values derived from the probability of noting S1 and vPM activations alone. In addition to these co-activation findings, we analyzed firing rates in a time-resolved fashion, which allowed us the opportunity to see whether firing rates to sensory stimulation during consciousness were more sustained than during unconsciousness. As predicted by the GNW and well established in electroencephalography and electrocorticography [54, 55, 56], neural activity was more sustained when animals were conscious (vs. unconscious). Taken together, our results provide empirical support (again through the lens of single neurons) for the GNW theory.

In addition to providing empirical neurophysiological evidence relevant to two prevailing theories of consciousness, our results make several novel contributions to the study of multisensory integration. First, to our knowledge, this is the first report to detail that supra-additivity and enhancement indices are normally distributed in both vPM and S1 of non-human primates (see [57], for a categorization of S1 neurons demonstrating enhancement and supra-additivity in rats). Second, we observed a large number of neurons exhibiting multisensory enhancement in S1 and vPM, yet very few that exhibited supra-additivity. These results comport well with the known multisensory convergence in vPM [58, 59], but also represent the first evidence that these neurons can integrate this information. Third, we detail the dispersion tendencies associated with the firing patterns of (multi)sensory neurons in S1 and vPM. Variance in neuronal firing may be a result of a variety of causes [33], both internal to the neuron or as a network property. Interestingly, while the Fano Factor is likely impacted by both these sources, an elevation in noise correlations likely reflects a source of co-modulation. Thus, the current results demonstrating an increase in both Fano Factor and noise correlations during unconsciousness suggests a dynamical system in which the firing patterns of individual neurons are becoming more chaotic yet the population as a whole is more synchronously co-activated; an observation that is in line with reports suggesting a potentiation of slow oscillations and a reduction of high-frequencies during unawareness [27]. Finally, from the observations that conscious states are seemingly associated with low noise correlations and that neurons showing multisensory convergence (as opposed to integration) more faithfully track consciousness according to this metric, we reasoned that perhaps a high degree of noise correlation is beneficial to multisensory integration. In fact, our results suggest a positive correlation between the amount a neuron shares noise with its neighbors, and the degree to which it exhibits multisensory integration. We find this result particularly interesting, as multisensory integration is a special form of integration – a form that has minimal shared variance at the periphery, since information is transduced by different sensory organs. This relationship between shared noise and greater multisensory integration may be a result of larger dendritic arborizations in integrative neurons (see [60, 61]), and may represent the neural instantiation of the postulation that stimulus correlation detection subserves the synthesis of information across the senses [62-64].

In conclusion, we started from the IIT [11] and the GNW [7, 8] to derive neurophysiological predictions relating to consciousness. We then leveraged multisensory neurons and circuits to functionally label neurons as convergent and integrative, and used these categorical distinctions to test IIT and GNW-derived predictions. The neurophysiological results generally support the GNW and not the IIT. However, important caveats exist. First, it is possible that the predictions we generated according to the IIT represented a higher bar to clear than those we generated from the GNW. Indeed, this is a strength of the IIT (i.e., making strong prediction) – and thus future work should aim at continuing to translate theoretical postulates into concrete hypotheses, and subsequently testing these hypotheses. Second, the IIT predictions derived and tested here represent the simplest implementation and interpretation of the theory possible. Only three nodes were used, and only a single node was changed in calculating different phi values. Further, we have assumed that single unit spiking activity was a good approximation of the behavior of “nodes” within the IIT. This is far from trivial, as for example, in the IIT formalism nodes are either “on” or “off”, yet real neurons can show graded levels of activity. Third, while simultaneously recording from S1 and vPM lends nicely to testing the GNW – given their known micro-circuitry [30, 31] – it may be argued that these areas are not ideal for testing the IIT. Indeed, Koch and colleagues [65], researchers supporting the IIT, recently suggest that anatomical correlates of consciousness are primarily localized to a “posterior hot zone”. Thus, in the future it may be interesting to test similar ideas to those presented here in the posterior parietal cortex (but see [66] for arguments suggesting that the neural correlates of consciousness are in the “front” of the brain). Beyond specific objections related to the IIT, it must be emphasized that the results reported here are exclusive for propofol anesthesia. Hence, generalization of these results to the broader domain of consciousness must be done with caution. Future work may aim at replicating the above-described findings during the administration of several distinct anesthetics – the union of effects safely being able to be ascribed to consciousness. Further, it must be acknowledged that IIT is primarily a theory of consciousness-level, while GNW primarily focuses of consciousness-content. These different aspects of consciousness are closely related (as one does not consciously perceive the external environment if in an unconscious state), but dissociable. Here both aspects were conflated (i.e., it is assumed animals did not hear the auditory tone during unconscious-level), as the primary aim was to contrast IIT and GNW from a neurophysiological perspective. However, in the future it will be interesting to dissociate these dimensions. Lastly, the degree to which these findings generalize from macaques to humans is unknown. Regardless of model organism (macaque, human, or other), we consider that using sensory stimulation from distinct and multiple sensory modalities (e.g., [44, 67, 68] – as highlighted in the current report - may afford important leverage in the study of perceptual awareness and consciousness level by allowing functional characterization of neurons and neuronal ensembles.

## Materials and Methods

### Animal Model

Animals were handled according to the institutional standards of the National Institutes of Health (NIH) and an approved protocol by the institutional animal care and use committee at the Massachusetts General Hospital. Two adult male monkeys (*Macaca mulatta*, 10 –12 kg) were used.

### Behavioral Task and Experimental Procedure

The animals were trained in a behavioral task wherein following the onset of a start tone (1000 Hz, 100 ms, see Figure 9A, first row) they were required to initiate each trial by holding down a button with their hand ipsilateral to the recording hemisphere. In order to successfully launch a trial (before loss of consciousness), the animals were required to hold the button within 1.5 seconds of the trial onset tone (Figure 9A, second row). Then, following button press, within a uniform random delay between 1 and 3 seconds (Figure 9, blue shaded area with dashed contour representing a variable delay) one of four sensory stimulus sets was delivered (tactile air puffs, T; auditory stimuli, A; simultaneous auditory and tactile, AT; no stimuli, N; Figure 9A depicts an AT trial, and hence T, A, and N trials are shaded). Air puffs during T trials were delivered at 12 psi to the lower part of the face contralateral to the recording hemisphere via a computer-controlled regulator with a solenoid valve (AirStim; San Diego Instruments). The eye area was avoided. Auditory stimuli during A trials were pure tones at 4000 Hz and at 80 dB SPL generated by a computer and delivered using two speakers 40 cm from the animal. Audiotactile (AT) trials were simply the joint and simultaneous presentation of A and T trials. N trials were catch trials were no stimulus was presented. White noise (50 dB SPL) was applied throughout the trial to mask inherent noise derived from air puff and mechanical apparatus. All of the stimulus sets were presented randomly to the animal regardless of their behavioral response throughout the recording session. Following the presentation of the sensory stimulus the animals were required to keep holding the button down until the presentation of liquid reward (3 seconds post stimuli onset, Figure 9A bottom row and second blue interval). The monkeys were trained to perform a correct response on >90% of the trials consistently for longer than 1.5 hours in an alert condition. The animal’s performance during the session was monitored and simultaneously recorded using a MATLAB-based behavior control system [69, 70]. Trial-by-trial behavioral responses were binned as a correct response (button holding until the trial end and release), failed attempt (early release, late touch, or no release of the button), or no response (Fig. 1C). Loss of consciousness was defined as the first no-response trial that was consistently followed by a lack of responses for the rest of anesthesia (see Figure 9B for an exemplar session where the cumulative sum of trials categorized as correct responses raises quickly initially and then saturates, while the cumulative sum of trials categorized as no-response is initially stagnant at zero and subsequently raises rapidly following approximately 280 trials).

**Figure 9.**
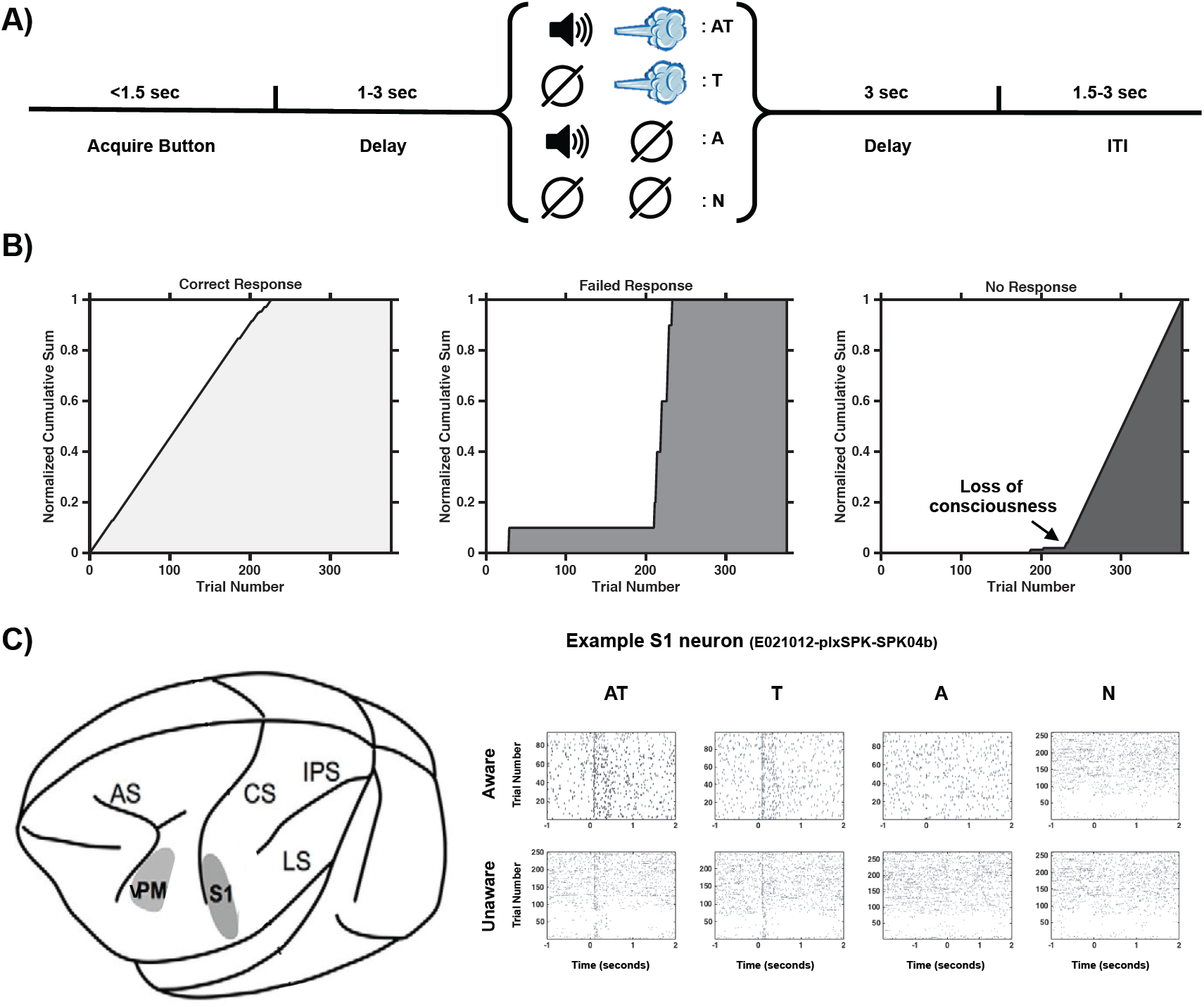
Experimental Procedure, Methods, and Neurophysiology Preprocessing. **A)** Experimental task; animals were required to press a button within 1.5 seconds following a start tone. Subsequently, following a random delay between 1 and 3 seconds post button press (dashed blue area) they were presented with a sensory stimulus (audiotactile, AT (purple); tactile, T (blue); or auditory, A (red)) or not (faded black, N). In this case an AT trial is illustrated, and hence represented in a continuous line, while T, A, and N are dashed and shaded. After a fixed delay of 3 seconds post stimulus onset, if the monkey was still holding the button, it was given a liquid reward and allowed to stop pressing the button. The trial depicted is a correct response trial, but a trial could also be categorized as failed response (e.g., released the button too soon) or a no-response trial (e.g., the monkey never executed button press). **B)** Cumulative sum of trial categories (leftmost; light gray = correct response; center, dark gray = failed response; rightmost, black = no response). Initially all trials are correct, but as propofol is administered, the animal falls unconscious and does not execute the button press. Unawareness is defined as the period between the first no-response trial that is consistently followed by a lack of responses for the rest of anesthesia. **C)** A schematic of a monkey brain depicting areas S1 and vPM, where neurons were recorded and example raster plots from a neuron in S1. Responses during an aware period are depicted on the top row, while the bottom row illustrates activity during unawareness. The first column shows audiotactile trials, the second illustrates tactile trials, the third shows audio trials, and the last column shows spiking activity during trials with no sensory stimuli. On the x-axis is time (in seconds, centered at stimuli onset) and on the y-axis is trial number.

### Anesthesia

Thirty minutes after initiating each recording session, propofol was infused for 60 minutes at a fixed rate (200 g/kg/min for Monkey E, and 230 or 270 g/kg/min for Monkey H) through a vascular access port. The infusion rate of propofol was a priori determined to induce loss of consciousness in approximately 10 minutes for each animal. No other sedatives or anesthetics were used during the experiment. The animal’s heart rate and oxygen saturation were monitored continuously throughout the session (CANL-425SV-A Pulse Oximeter; Med Associates). The animals maintained >94% oxygen saturation throughout the experiments.

### Neurophysiology Data Recording and Preprocessing

Before starting the study, a titanium head post was surgically implanted on each of the two animals. A vascular access port was equally surgically implanted in the internal jugular vein (Model CP6; Access Technologies). Once the animals had mastered the behavioral task described above, extracellular microelectrode arrays (Floating Microelectrode Arrays; MicroProbes) were implanted into S1 and vPM through a craniotomy (see Figure 9C). Microelectrodes were also implanted in S2, but due to insufficient recorded neurons caused by a technical malfunction, here we focus our report on recordings from S1 and vPM. Each array (1.95×2.50 mm) contained 16 platinum–iridium recording microelectrodes (0.5 MΩ, 1.5– 4.5 mm staggered length) separated by 400 µm. Landmarks on cortical surface and stereotaxic coordinates [71] guided the placement of arrays. A total of five arrays were implanted in Monkey E (two arrays in S1, one in S2, and two in vPM, all in the left hemisphere) and four arrays in Monkey H (two arrays in S1, one in S2, and one and vPM; all in the right hemisphere). The recording experiments were performed after 2 weeks of recovery following the array surgery. All experiments were conducted in a radio frequency shielded recording enclosure.

Neural activity was recorded continuously and simultaneously from S1 and vPM through the microelectrode arrays while the animals were performing the behavioral task. Analog data were amplified, band-pass filtered between 0.5 and 8 kHz, and sampled at 40 kHz (OmniPlex; Plexon). The spiking activity (see Figure (9C) was obtained by high-pass filtering at 300 kHz and applying a minimum threshold of 3 standard deviations in order to exclude background noise from the raw voltage tracings on each channel. Subsequently all action potentials were sorted using waveform principal component analysis (Offline Sorter; Plexon) and binned into 1 ms bins, effectively rendering the sampling rate 1kHz.

### Neurophysiology Data Analyses

#### Firing Rate and Fano Factor

Both central and dispersion tendencies of single-unit spiking activity in S1 and vPM were quantified as a function of stimulus modality as it is well-established that mean firing rates alone do not fully characterize the properties of neural activity [33]. Regarding firing rates, spikes were first binned in 1ms intervals, and epochs were centered on stimuli onset, ranging from 2000ms prior to stimuli onset (i.e., −2000ms), to 2000ms after stimuli onset. Subsequently spike counts were effectuated within a 100ms window, between −500ms and 1000ms, and in steps of 10ms. It must be noted that this analysis essentially low-passes time, and hence the exact timing of reported effects should not be emphasized. Analyses of firing rates were conducted both on baseline-corrected and non-corrected rates. The contrast of non-corrected rates allows for determining the impact of propofol on baseline firing, while the analysis on baseline-corrected rates allows specifically querying the evoked-responses to stimuli onset. That is, for the baseline-corrected rates, every spike count function was centered along the y-axis (i.e., spikes/s) to zero according to their own baseline firing (−500 to 0ms post-stimuli onset). In this manner, positive deviations from 0 indicate an increased in firing rate, while negative deflections indicate a silencing in spiking activity post-stimuli onset with respect to baseline. Spike counts were first averaged within a cell and across trials, and subsequently across neurons. In terms of statistical analyses, as the temporal dynamics of spiking activity was of interest, in particular within the GNW theory [7, 8] emphasizing sustained activity in aware and not unaware states, we conducted a time-resolved (at each 10ms time-point, 151 in total) 2 (State; Aware vs. Unaware) × 2 (Area; S1 vs. vPM) × 4 (Stimulation; AT, T, A, N) independent samples analysis of variance (ANOVA). As spiking rates were not normally distributed (i.e., presence of a true floor, in that negative spikes are not possible), the ANOVAs for non-baseline corrected rates were conducted on log-transformed data. On the other hand, the subtraction of evoked activity to baseline activity did yield normal distributions, and hence this data is analyzed without log-transform. The inter-dependence of observations is difficult to ascertain within a neural network composed of neurons whose precise connections are unknown, and thus independent as opposed to dependent ANOVAs were conducted in order to adopt the most conservative approach (i.e., within-samples ANOVAs are statistically stronger than between-samples analyses). Similarly, in order to protect against Type I error (i.e., false positives) significant effects were only considered at α < 0.01 for at least 3 consecutive windows (i.e., 30 time-points; see [44], for a similar approach with time-series data).

Regarding the inter-trial variance in evoked responses associated with the distinct states of consciousness, stimuli modalities, and brain areas, fano factors (i.e., ratio of spike-count variance to spike-count mean) were calculated [70]. Indeed, repeated trials do not yield identical responses, and this variance is associated both with cellular and molecular processes involved in spike generation at the axon hillock (e.g., refractory periods) and network-level properties [63]. Conveniently the neuron-specific variance is largely consiered to be well accounted by a Poisson point process (i.e., mean and variance scale), and hence a fano factor of 1 [33, 64]. Fano factors in excess of 1, thus, may be considered to index variability that is associated with network-level properties and this variability is typically reduced at stimuli onset. Here, therefore, we report time-resolved fano factor both corrected for baseline (in order to examine putative network-level decreases in inter-trial variability as a function of stimuli onset, awareness state and sensory modality), and not corrected for baseline (in order to assess basal cell-specific and network level inter-trial variability as a function of awareness state). Statistical analysis is conducted as described above for firing rates.

#### Neural Index of Multisensory Integration

The hallmark for multisensory integration at the single unit level is an evoked response to multisensory stimuli (e.g., AT) that may not be linearly predicted by responses to the constituent unisensory stimuli (e.g., A and T; [73]. Thus, given the time-resolved results demonstrating sustained activity to sensory stimulation until approximately 500ms post-stimuli onset, mean spike counts to AT, T, A, and N trials were executed within this time period (see [34], for a similar approach). Subsequently, the i) supra-additivity and ii) enhancement index of each neuron was computed (according to Eq. 1 and Eq. 2, respectively). Historically, supra-additivity – the degree to which a multisensory response exceeds the sum of unisensory responses (Eq. 1) - was considered the clearest indication of multisensory facilitation); nonetheless this feature is not as prominent in cortex as it is in sub-cortex [39, 73]. Thus, we supplement the supra-additivity index with the enhancement index – the degree to which a multisensory response is greater than the maximal response to unisensory stimuli (see Eq. 2). An enhancement index above 1 indicates a neuron that is further driven by multisensory than unisensory stimulation. Supra-additivity (Eq. 1) and enhancement (Eq. 2) indices were computed as follows;

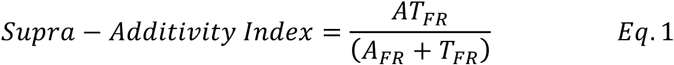

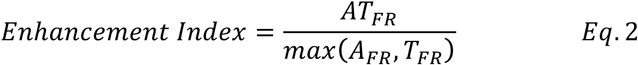

where *AT*_*FR*_ is the mean baseline-corrected firing rate for a particular neuron to audiotactile stimulation, *T*_*FR*_ is the mean baseline-corrected firing rate for the particular neuron to tactile stimulation, and finally *A*_*FR*_ is the mean baseline-corrected firing rate for the particular neuron to auditory stimulation.

#### Bifurcation Into Convergence and Integration

Modeling results based on the IIT specify that a network converging on a neuron that integrates information, as opposed to responding indiscriminately, ought to support a greater degree to consciousness. Hence, here we aim at testing two predictions that may follow from the IIT; i) as an organism falls into unconsciousness, the neurons that are most impacted are those that integrate information (i.e., putatively anesthetics act on these neurons preferentially), and ii) neurons that integrate information exhibit the properties of consciousness when the organism is conscious. To test these predictions, we divide our population of neurons into those that integrate vs. converge (Figure 4 and beyond). However, initially we simply describe the proportion of neurons that fit within each category (Figure 5) in a non-mutually exclusive fashion. A neuron that converges information is defined as a neuron that on average (i.e., across trials) responds – spike count from 0 to 500ms - to both unisensory auditory and tactile information beyond its baseline firing rate (−500ms to 0ms) plus 2 standard deviations. That is, in order to qualify as convergent, the spiking count of a neuron to AT stimulation does not need to be examined. On the other hand, a neuron that integrates information is defined as a neuron that is most readily driven by the simultaneous presence of A and T information. Thus, neurons that respond to AT stimulation (as defined above) and do so to a greater degree than their maximal unisensory response (i.e., enhancement index above 1) were initially classified as integrative. Importantly, beyond Figure 3 (e.g., to categorize the fate of neurons when the animal becomes unconscious and quantify neural complexity, noise correlations, and neural ignition) two mutually exclusive classes are created. Neurons that respond indiscriminately to sensory stimulation and not preferentially to multisensory vs. unisensory presentations are classified as convergent, while those that exhibit multisensory enhancement without being considered convergent are taken to integrate information. Given the initial number of neurons in S1 and vPM, this bifurcation yielded a sufficient quantity of neurons exclusively categorized as convergent (N = 125) and integrative (N = 64) in S1, but not in vPM (convergent, N = 61; integrative, N = 8) – thus, for the analyses specifically probing the difference between convergent and integrative neurons, analyses are restricted to S1. Further, given the heterogeneity of neuron’s spike trains (see Figures 1-3) for Figure 3 and beyond we considered a neuron as fitting within a particular category (e.g., A, T, AT convergent, AT integrative) if at some point between 0ms and 1000ms post-stimuli onset they met the particular criteria for at least 50 consecutive ms.

Equally of note, in Figure 3 neurons that are labeled to integrate auditory and tactile information (purple, orange, and green) are not first indexed for their unisensory responses. That is, while in S1, 49% of neurons are classified as responding to a greater extent to AT stimulation than to the maximum unisensory stimulation, this latter unisensory response is not necessarily different from baseline activity. We consider this approach appropriate within the current aim of leveraging multisensory responses in querying consciousness theories, but it must be highlighted that multisensory enhancement may be more strictly considered to apply only when tactile, auditory, and audiotactile responses are different from baseline, and the latter responses is greater than the maximal of the former two [27]. Indeed, the categorization here is more in line with the recent emphasis within the study of multisensory integration to index covert multisensory processes [36], in particular within classically considered primary sensory areas [74], than with the original description of multisensory integration in the late eighteens and early nineties [73].

#### Lempel-Ziv Complexity

Categorizing the complexity of neural representations – operationalized as the number of distinct patterns present in data – has become of increasing popularity as of late (e.g., [11; 33]) in particular due to its ability to differentiate between states of consciousness given scalp electrophysiological data [6] and the belief that complexity is at least indirectly related to functional differentiation/integration, paramount notions with the IIT [11]. In order to quantify neural complexity, here we measure the Lempel-Ziv (LZ) complexity [28] associated with each spike train evoked as a consequence of AT, T, A, or N trials, and as a function of the animals’ consciousness state. LZ complexity measures the approximate quantity of non-redundant information contained within a string by estimating the minimal size of the “vocabulary” necessary to describe the entirety of the information contained within the string in a lossless manner [28]. That is, it is a lossless compression algorithm (routinely used in ZIP files and TIFF images), and it is utilized to measure the number of distinct patterns in symbolic sequences, in particular within binary signals. LZ is impacted by the overall entropy within a signal [45]; i.e., a binary string composed almost exclusively of ‘0’ will not have a high LZ, not due to the arrangement of those ‘1’, but simply because there are not many of them). Thus, here, to equate entropy across conditions we first converted spike trains into a continuous measure by convolving each trial with a Gaussian kernel with σ=50ms, and then binarized each time-point within this trial by assigning a ‘1’ to time-points above the trial mean, and ‘0’ to time-points below the trial mean. Next, LZ was computed (28) in MATLAB within a sliding window moving between −500ms and 750ms post-stimuli onset, a length of 100ms, and step size of 50ms. Lastly, the same procedure was executed while randomly shuffling the binary sequence before calculating LZ. This shuffled LZ time-series represents a theoretical upper bound (i.e., random data has a higher LZ) and was used to normalize the calculated LZ from the non-shuffled data. Hence, a normalized LZ of 1 indicates ‘as complex as random noise’, while lower values indicate the presence of structure in the data (see [43, 44], for a similar approach). Statistical analysis largely followed that of firing rates and fano factors, which exception that data were never log-transformed as they were normally distributed. Analysis was effectuated both on baseline-corrected values, in order to compare the negative deflection present during stimulus onset (see [43] for a similar findings) and most importantly, on non-corrected values, in order to examine the basal complexity in spiking activity as a function of consciousness and whether neurons were categorized as convergent or integrative.

#### Noise Correlations

While LZ complexity is arguably the most often utilized measure within the IIT framework [11], it is not a traditional measure within neurophysiology. Thus, we sought to further probe the properties of convergent and integrative neurons – and their correspondence with the alteration in the particular measure as a function of consciousness state – with a neurophysiological measure that is well established to alter with consciousness state. Noise correlations [29] express the amount of covariability in the trial-to-trial fluctuations of responses of two neurons to repeated presentations of the same stimuli, are central to questions of coding accuracy and efficiency [75], and are well-established to be altered by consciousness state [29]. Thus, this measure was computed both in S1 and vPM neurons, as a function of consciousness state and stimuli modality. Noise correlations where computed as the Pearson correlation between all pairs of neurons recorded simultaneously within the same session (see [29] for a similar approach). Spike counts were effectuated for each trial on the 500ms immediately following stimuli presentation (defined above as the average time-period of neural response, and in concert with [29]. We considered the noise correlation for a particular neuron it’s average correlation with all other neurons recorded in the same session.

#### Neural Ignition

The GNW model points to the late amplification of relevant sensory activity, long-distance cortico-cortical synchronization at beta and gamma frequencies, and ignition of large-scale fronto-parietal networks as neural measures of consciousness [8]. To test this prediction, we query at the single trial level whether sensory stimulation leads to co-activation of both primary sensory areas (i.e., S1) and frontal regions (i.e., vPM) more commonly during conscious than unconscious states. For each neuron (both in S1 and vPM) we specify a threshold benchmarking reliable neural activity as the average spike count between −500 and 0 ms post-stimuli onset plus 2 standard deviations. Similarly, the neural response is considered to be the spike-count between 0 and 500 post-stimuli onset. Then, iteratively we pick a neuron from S1 and a neuron from vPM and query whether on a particular trial did neither area respond, did solely S1 respond, did solely vPM respond, or did both S1 and vPM respond. A particular S1 neuron is subsequently paired with all neurons in vPM recorded during the same session, and finally it’s mean activation patterns (e.g., S1 and vPM active, vPM active, S1 active, or none) as a function of consciousness state and sensory stimulation are quantified. The same procedure is applied to vPM neurons. It must be highlighted that routinely mean firing rates are largely driven by strong responses in a few trials [33], for example), and hence demanding a response within a particular trial to exceed baseline plus 2 standard deviations is a conservative approach yielding a great number of no-response trials. Nonparametric statistics are used in this analysis as data did not confirm to the assumptions made by parametric inference statistics.

## Acknowledgements

JPN was supported by F31MH112336. The authors thank Dr. Randolph Blake for insightful comments on an earlier version of this manuscript.

## Author Contributions

JPN, EE and MW conceived of the experiment. YI and SP collected all experimental data. JPN performed the computational work. YI and SP performed data pre-processing and JPN analyzed the data. JPN drafted the initial version of the manuscript. All authors edited the manuscript and approved of the final version.

## Declaration of Interests

The authors declare no competing interests.

## SUPPLEMENTARY INFORMATION

### Rationale and Computation of Integrated Information (Φ)

From an information-theoretic perspective information is the reduction of uncertainty (Shannon, 1948). In turn, information may be quantified by considering how a system in its current state *S*_0_ constrains the system’s potential past and future states. Figure S1 illustrates this principle form within the purview of *C* at time *t* for the system with an XOR gate. Under this scenario, if *C* is currently active, then at time *t*-1 by necessity either A was active, B was active, A and C were active, or B and C were active (Figure S1, left panel). The probability distribution of past states that could have been causes of *C* = 1 is its cause repertoire *p ABC*^*past*^ |*C* = 1. On the other hand, if it is unknown in what state *C* is in, *t*-1 is unconstrained *p*^*uc*^(*ABC*^*past*^). A similar rationale applies to future states wherein the current state of *C* constrains its future potential states, and the effect repertoire is thus the probability of being in any given state given that *C* is current active, or *p ABC*^*future*^ |*C* = 1. The amount of information that *C* = 1 specifies about the past is its cause information (CI) and the amount it specifies about the future is its effect information (EI). CI and EI are respectively measured as follows,

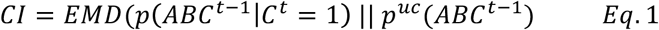

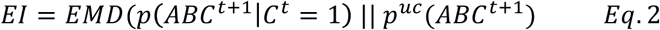

where *EMD* refers to earth mover’s distance (Rubner et al., 2000), the minimal cost of reshaping one distribution (e.g., unconstrained) into the other (e.g., constrained) or area of distribution moved times the distance moved. Finally, the total amount of cause-effect information (CEI) specified by *C* = 1 is the minimum value between CI and EI. This results from the fact that both CI and EI may act as limiting cases – an information bottleneck – and hence minimize the CEI of the system as a whole (see Oizumi et al., 2014, for detail). Finally, while CEI measures information, the IIT conjectures that consciousness is *integrated* information. That is, information generated by the system above and beyond that generated by its constituent parts. Hence, the system as a whole is iteratively partitioned into all possible subsystems or purviews and the process delineated above is evaluated for each of these components. Similar to CEI, integrated information is calculated as the *EMD* between the cause-effect repertoire specified by the system as a whole and the cause-effect repertoire of the partitioned system. Φ is the distance between the system as a whole and the system-partitioned that makes the least difference; the minimum information partition. That is, Φ is the degree to which the cause effect repertoire for the system as a whole differs from the next most informative partition.

**Figure S1.**
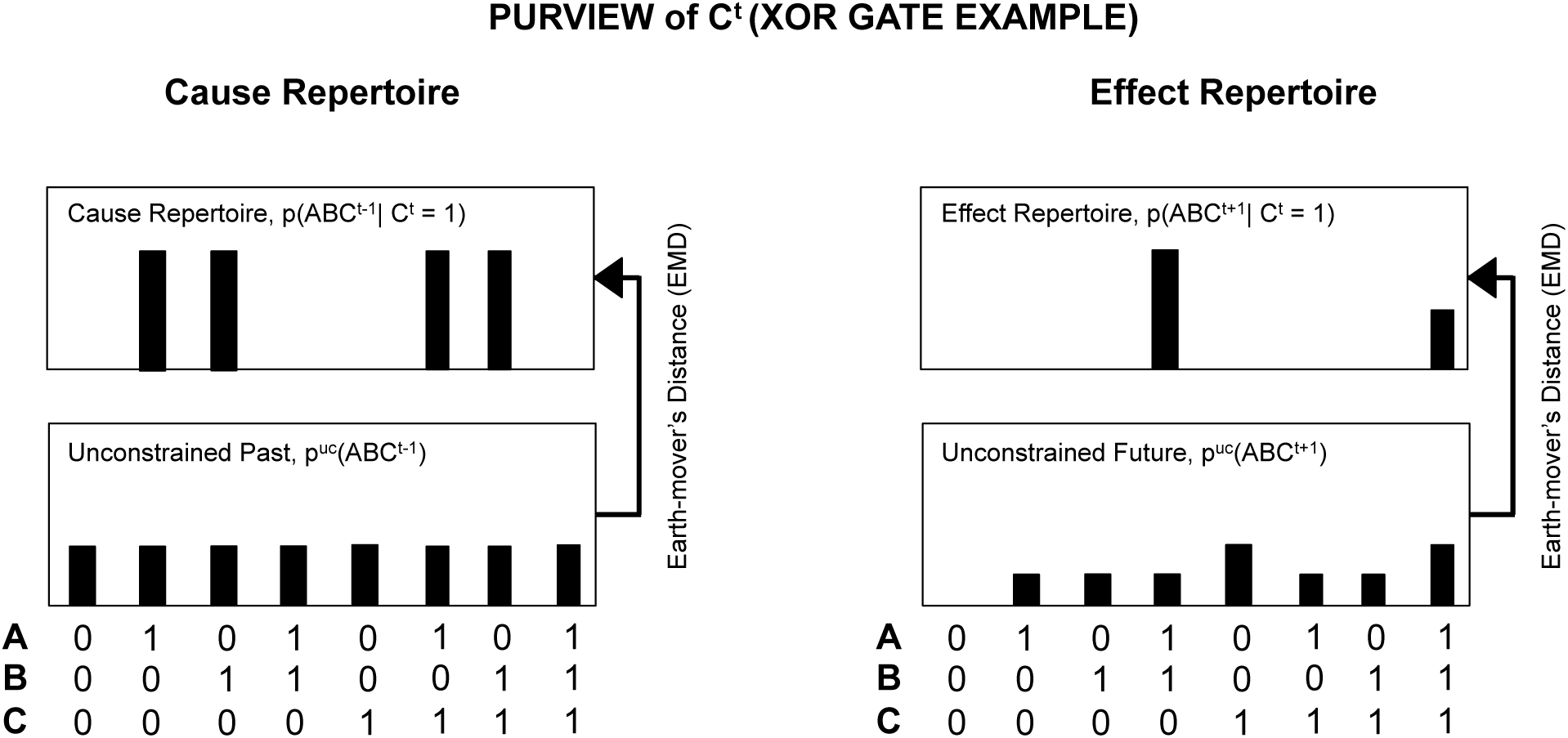
Illustration of cause and effect repertoires and the constraints imposed on potential probability distributions by the fact that C=1. Cause (left) and effect (right) repertoires for a system with three nodes as the one illustrated in Figure 1, and as a function of whether the past-future is constrained to C=1 (top) or not (bottom).

Information integration (phi, Φ) was calculated for the multisensory convergent and integrative networks using the transitions probability matrices illustrated in Figure S2 (see below) and as implemented in *PyPhi* (Mayner et al., 2017) with Python 3.4.

### Formalizing the Role of Multisensory Integration in Consciousness

To formally ascertain the putative role multisensory integrative (vs. convergent) neurons within a network in bearing consciousness (according to the IIT), we built two biologically inspired simple neural networks (Figure S2A). These networks each have 3 nodes, two of which may be considered analogous to unisensory areas (nodes A and B) and a third (node C), which receives projections from the unisensory areas and may be considered analogous to a multisensory area. As is well established in biological systems, the multisensory area equally projected back to unisensory areas (Bizley et al., 2007, Cappe et al., 2009, Ghazanfar and Schroeder, 2006). The two networks were identical with exception that for one network (Figure S2A, left panel) node C was an “XOR” gate, while for the other it was an “AND” gate (Figure S2A, right panel). The XOR gate results in a logical “true” (or ‘1’/ ‘HIGH’) when the number of true inputs is odd. In this case, given the system architecture, the node C would be active if on the previous time step one and only one of gates A or B was active. Thus, node C can in principle be active following activity in either node A or B, but importantly does not respond preferentially when both are active. On the other hand, the other network, where node C is an “AND” gate, responds exclusively when on the precedent time-step both gates A and B were active. That is, this gate most faithfully mimics the behavior of integrative multisensory neurons that may or may not overtly respond indiscriminately to distinct sensory inputs, but importantly are most driven by spatio-temporally coincident multisensory inputs. Hence, the network formed with an XOR gate (Figure S2A, left) instantiates a network with neurons that are indiscriminant to the nature of sensory input, while in contrast the network formed with an AND gate (Figure S2A, right) instantiates a network with neurons that integrate sensory information, i.e., responds non-linearly to the addition of sensory stimuli from distinct modalities (Stein and Stanford, 2008, Wallace et al., 2006). The architecture of these systems dictates the composition of transition probability matrices (TPMs), which guides transitions between states (i.e., neurons that are ‘active’ at different time-points). In Figure S2B these TPMs have been depicted (left and right respectively for the multisensory convergent and multisensory integrative systems) and highlighted for their distinctive features. Namely, in the case of the convergent network, when ABC nodes are in state 100 (respectively, A, B, and C) or 010 (green rows), activation of the multisensory node will follow. This is not true if the convergent network is in state 110 (red row). The opposite is true for the network that integrates. Given these TPMs, Φ can be calculated when the state of the network is ABC = 001 (multisensory node active). Results indicate that in fact a network with a node with integrative capacity in principle may bear a greater degree of consciousness (Φ = 0.78) than one that simply responds indiscriminately to stimuli from distinct sensory modalities (Φ = 0.25).

**Figure S2.**
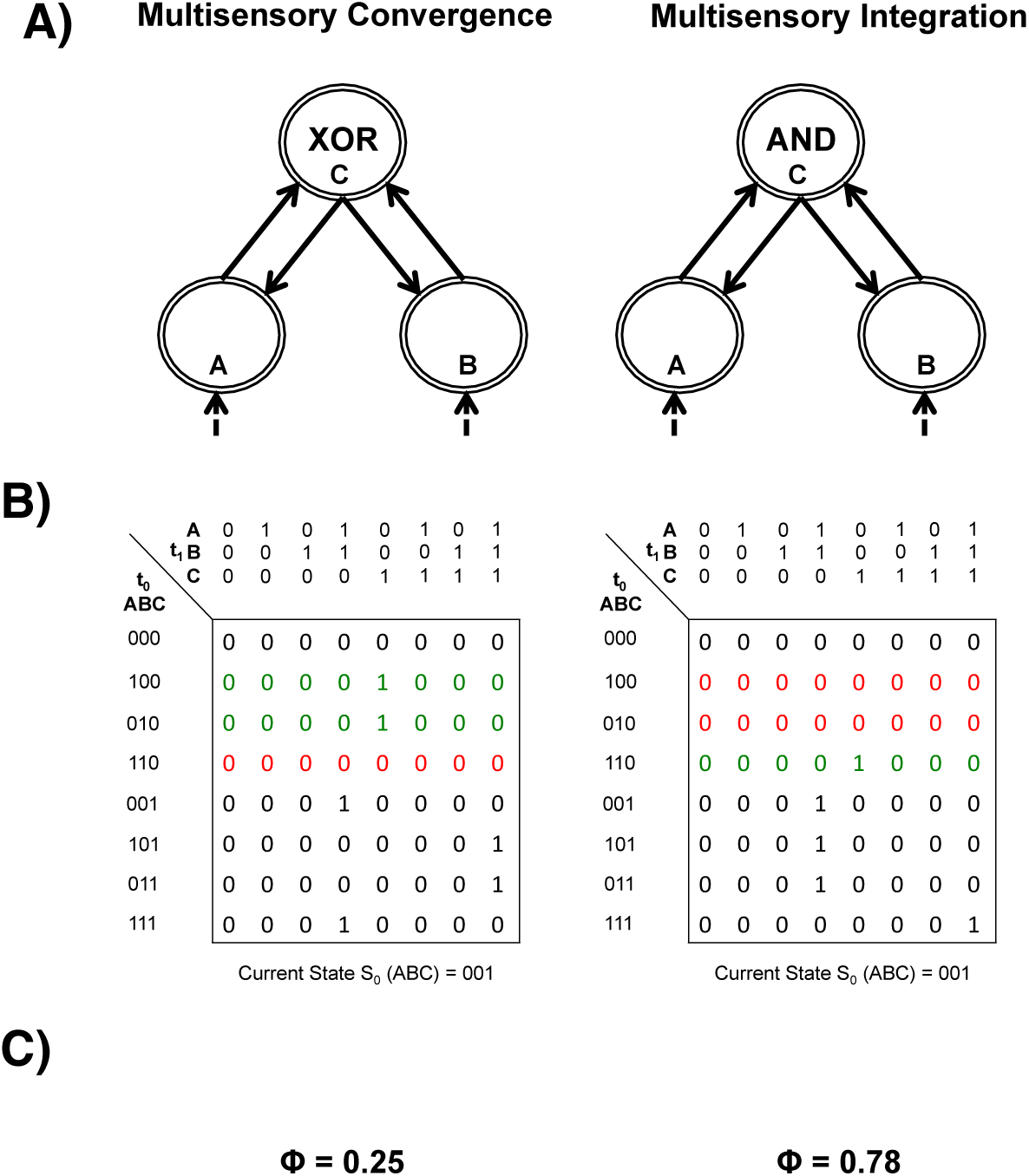
Formalizing the role of multisensory integrative neurons in bearing consciousness according to IIT. **A)** Depiction of a multisensory convergent (left) and integrative (right) network. There is no connection between A and B nodes, as these transition probability values are zero. The dashed connections leading to A and B are to illustrate that these putative unisensory areas receive input from downstream neural areas, yet they play no role in the ITT-driven model. **B)** The transition probability matrices (TPM) for a deterministic (e.g., probability is either 0 or 1) convergent (left) and integrative (right) network are illustrated. State t is represented in the abscissa and t+1 on the ordinate. Green and red rows are highlighted to emphasize key difference between the convergent and integrative networks, yet these differences are not exhaustive (however do dictate the rest of differences). **C)** The Φ value associated with the convergent (left) and integrative (right) TMPs as determined in *PyPhi* (Mayner et al., 2017).

### S1 and vPM Firing Rates as a Function of Sensory Modality and Awareness

Regarding the firing rate, analyses on non baseline-corrected activity indicated a clear generalized decrease in firing rate when monkeys were rendered unconscious (p<0.01 at all time points; Aware; M = 4.43 spikes/s, S.E.M = 0.008 spikes/s; Unaware; M = 2.44 spikes/s, S.E.M = 0.007), a significant difference in spiking activity across the areas recorded between 50ms and 160ms post-stimuli onset (p<0.01, S1, M = 5.68 spikes/s, S.E.M = 0.008 spikes/s; vPM, M = 4.88 spikes/s, S.E.M = 0.006 spikes/s), and a significant main effect of stimulation type (i.e., AT, T, A, N) between 50ms and 480ms post-stimuli onset (AT, M = 4.14 spikes/s, S.E.M = 0.01 spikes/s; T, M = 4.31 spikes/s, S.E.M = 0.01 spikes/s; A, M = 3.75 spikes/s, S.E.M = 0.006 spikes/s; N, M = 3.28 spikes/s, S.E.M = 0.001 spikes/s). Importantly, a stimulation-type (i.e., AT, T, A, N) by area recorded (i.e., S1 vs. vPM) interaction was also evident (p<0.01, between 60ms and 210ms post-stimuli onset), driven by the fact that vPM responded to A stimulation (M = 3.21 spikes/s, S.E.M = 0.10 spikes/s), while S1 did not (M = 2.18 spikes/s, S.E.M = 0.10 spikes/s). Thus, in sum and as expected, these analyses demonstrated that propofol silenced spiking activity generally, that neurons in S1 and vPM responded differently to distinct sensory stimuli between 50 and 480ms post-stimuli onset, and that vPM but not S1 responded to auditory stimulation. The baseline-corrected analyses (depicted in Figure 2, rows 1 and 3) largely demonstrated analogous results, while indicating that the bifurcation in evoked responses (as opposed to baseline responses, as indicated above) between states of consciousness occurred (regardless of sensory modality) 80 ms post-stimuli onset (p<0.01, averaged across 80-1000ms post-stimuli onset; Aware, M = 0.48 spikes/s, S.E.M = 0.04; Unaware, M = 0.09 spikes/s, S.E.M = 0.01 spikes/s) and also highlighting a consciousness state by sensory modality (p<0.01 between 40-110ms, and 200-380ms) as well as 3-way (modality, state, and area) interaction (p<0.01, 410-610ms post-stimuli onset). The time-periods demonstrating a significant difference in evoked activity as a function of state of consciousness are shaded in gray in Figure 1 (main text) separated by area recorded and sensory stimulation, while the time-periods demonstrating a significant response vis-à-vis baseline are indicated by horizontal lines in each panel (see Figure 1).

### S1 and vPM Fano Factors as a Function of Sensory Modality and Awareness

Regarding fano factors (FFs), results demonstrated heightened variability under unaware (M = 1.45, S.E.M = 7.3e-4) than aware (M = 1.16, S.E.M = 5.6e-4) conditions (see Ecker et al., 2014 for a similar result), while both of these were significantly greater than 1 (unaware, p < 4.76e-92; aware, p = 4.76e-92), and hence likely exhibiting inter-trial variability above and beyond what is presumed to be attributable intrinsically to neurons (i.e., FF = 1). Similarly, FFs were larger in S1 (M = 1.32, S.E.M = 6.91e-4) than vPM (M = 1.22, S.E.M = 5.70e- 4), throughout the post-stimuli period (p<0.01, for exemption of the period between 80ms and 120ms post-stimuli onset. The period between 50ms and 270ms post-stimuli onset demonstrated a significant difference in FFs as a function of stimulus modality (p<0.01), with the AT (M = 1.29, S.E.M = 0.02) and T (M = 1.28, S.E.M = 0.03) conditions being the less variable (AT vs. T, p = 0.58) than the A (M = 1.31, S.E.M = 0.02) and N (M = 1.33, S.E.M = 0.02) conditions (T vs. A, t = 2.03, p = 0.041; A vs. N, p = 0.43). Importantly, FFs also demonstrated a consciousness state by recording area interaction (p<0.01, between 200ms and 320ms post-stimuli onset) and a recording area by stimulus modality interaction (p<0.01, between 50ms and 180ms post-stimuli onset). The latter interaction was driven by a main effect of stimuli modality that was sustained in S1 (p<0.01, between 50ms and 250ms, as well as 350ms and 540ms post-stimuli onset) and only transient in vPM (p<0.01, between 110 and 140ms post-stimuli onset), while the former is attributable to a rapprochement in FF between consciousness states in S1 and not in vPM. Indeed, this last effect is further appreciable when correcting FFs for baseline (Figure 1) as a further quenching in variability in S1 (vs. vPM) specifically to AT and T sensory stimulation (contrasts between aware and unaware conditions; S1; AT, p<0.01 between 160ms-200ms, T, p<0.01 between 180ms-220ms, never for A and N; vPM, never). As for firing rates, the time-periods demonstrating a significant difference in FF as a function of state of consciousness are shaded in gray in Figure 1 (main text) separated by area of recording and sensory stimulation type. Time-periods demonstrating a significant quenching in FF vis-à-vis baseline are indicated by horizontal lines in each panel (see Figure 1).

### Lempel-Ziv Complexity as a Function of Sensory Modality and Awareness

Figure 5A illustrates normalized LZ (see Andrillon et al., 2016, Noel et al., 2018, and Methods), both in its baseline-corrected and non-corrected format, and as a function of consciousness state (aware = colored; unaware = gray) and sensory stimulation. Regarding the non-corrected values, a 2 (consciousness state; aware vs. unaware) × 2 (recording area; S1 vs. vPM) × 4 (stimulation type; AT, T, A, N) ANOVA most strikingly revealed that unaware states (M = 0.88, S.E.M = 0.003) were generally more complex (p<0.01 at all time-points) than aware states (M = 0.81, S.E.M = 0.004). This analysis also revealed a main effect of recording area between 50ms and 100ms post-stimuli onset (p<0.01), as well as a main effect of stimulation between 50ms and 250ms (p<0.01). This analysis equally indicated a significant interaction between recording area and stimulation type (p<0.01 between 50ms and 150ms post-stimuli onset). The interaction was driven by a significant main effect of stimulation that lasted longer (p<0.01, between 50ms and 250ms post-stimuli onset) in S1 than vPM (p<0.01 between 100 and 150ms). Once normalized LZ was corrected for baseline, analyses specified a main effect of consciousness state specifically between 200 and 400ms post-stimuli onset (p<0.01), indicating that not only was overall LZ different across consciousness states, but the evoked nature of this measure equally differed. This main effect was driven by the AT and T conditions, where complexity returned to it’s baseline value more readily under unaware (AT, and T, return to baseline at 300ms) than aware states (AT and T, return to baseline at 350ms). The rest of statistical contrasts followed the same pattern as for the non-corrected values. The time-periods demonstrating a significant difference in evoked activity as a function of state of consciousness are shaded in gray in Figure 5A separated by area recorded and sensory stimulation, while the time-periods demonstrating a significant response vis-à-vis baseline are indicated by horizontal lines in each panel (see Figure 5A). In sum, therefore, the state of awareness is seemingly indexed in spiking activity by an overall lower level of LZ complexity (see Figure 5A, non-corrected normalized LZ), as well as by a more sustained negative deflection evoked by sensory stimulation (see Andrillon et al., 2016, for a similar observation).

### Percentage of Trials Evoking Significant Firing in S1 and vPM as a Function of Sensory Modality and Awareness

The percentage of trials that resulted in significant firing of S1 neuron was altered by the sensory modality of the stimuli presented and the consciousness level of the monkey. In particular, results demonstrated a main effect of consciousness state (Z=1294, p<0.001), stimulation modality (χ^2^=51.52, p<0.001), and an interaction between these variables (χ^2^=80.99, p<0.001). The interaction was driven by a significant main effect of stimuli type during consciousness (χ^2^ =91.18, p<0.001), but not unconsciousness (χ^2^=4.07, p=0.19). Regarding significant activation of pre-frontal cortex neurons, once again results demonstrated further activation consciously (M=6.9%) than unconsciously (M=3.1%; Z=1319, p<0.001), a main effect of stimulation type (χ^2^=105.7, p<0.001), and an interaction between these variables (χ^2^ = 233.11, p < 0.001). The interaction was driven by differential trial-activation percentages as a function of stimulation type when the monkeys were conscious (χ^2^ =133.7, p<0.001) but not unconscious (χ^2^= 7.51, p=0.08).

